# Multi-omic analysis reveals nitric oxide dependent remodeling in classically activated macrophages and identifies negative regulation mediated by AKR1A1

**DOI:** 10.64898/2025.12.31.697202

**Authors:** Nicholas L. Arp, Uzziah S. Urquiza, Marcel Morgenstern, Jonathan H. Schrope, James A. Votava, Steven V. John, Jack J. Stevens, Anna Huttenlocher, Joshua J. Coon, Jing Fan

## Abstract

Nitric oxide (NO•) is an important signaling molecule in many biological processes, including immune response. During response to classical activation stimuli lipopolysaccharide (LPS) and interferon-γ (IFNγ), macrophages generate NO• via inducible nitric oxide synthase (iNOS). To comprehensively define the effects of NO•, we applied a multi-omic strategy integrating proteomics and transcriptomics to profile murine macrophages across conditions with or without LPS/IFNγ-activation, with or without iNOS expression or exogenous NO• donor treatment. The results revealed NO• has broad, yet selected and controlled, regulatory effects, playing a key role in coordinating the systematic remodeling during macrophage classical activation. Among the proteins that are most suppressed in a NO•-dependent manner, electron transport chain (ETC) is the most enriched. NO• drives complex-specific remodeling of ETC, causing selected downregulation of complex I, II, and IV, through a different combination of transcriptional and post-transcriptional mechanisms for each complex. Functionally, we found NO• is required, but not sufficient, for the strong suppression of cellular respiration upon macrophage activation. Among the most consistently upregulated proteins are many enzymes involved in redox defense. AKR1A1 was identified as a top hit. We found *Akr1a1* induction requires both NO• and LPS/IFNγ stimulation. The S-nitroso-CoA reductase activity of AKR1A1 mitigates NO•-driven inhibition of pyruvate dehydrogenase complex by limiting the inhibitory modifications targeting its lipoyl cofactor. Knocking out *Akr1a1* causes accelerated remodeling of TCA cycle, dysregulated immunoregulatory metabolite level, and altered functional gene expression and cytokine production at later stage of immune response. Thus, the NO•-dependent upregulation of AKR1A1 forms a negative regulatory loop to fine-tune NO•-mediated metabolic and functional remodeling during immune response. Together, this work provided a systems-level map of NO•-dependent regulation, revealed the crosstalk between NO• and immune signaling, and demonstrated mechanisms providing redox adaptation and precise control of NO•’s effects.

## INTRODUCTION

Nitric oxide (NO•) is a crucial signaling molecule that regulates diverse physiological processes across biological systems. Through mechanisms including regulating transcriptional programs, post-translational modification of proteins, binding to effector proteins and altering metal centers, and effects on low-molecular weight redox metabolites such as coenzyme A (CoA) and glutathione, NO• influences protein activity, cellular redox state, cellular metabolism and functions [1]. These coordinate processes ranging from vasodilation and neurotransmission to pathogen control and immune responses [2,3]. NO• functions both as a signaling mediator and as a metabolic effector. Although individual NO•-sensitive proteins have been extensively characterized, how NO• shapes cellular regulatory programs at a systems level remains incompletely understood.

Macrophages represent a biologically relevant model for examining NO•-dependent regulation [4]. During classical activation with lipopolysaccharide (LPS) and interferon-γ (IFNγ), macrophages produce NO• via inducible nitric oxide synthase (iNOS). The NO• production generates a reactive nitrogen species (RNS)-rich environment and temporally correlates with broad cellular remodeling [5]. The cellular remodeling upon classical activation encompasses transcriptional, proteomic, redox, and metabolic changes [4,6–13]. However, which parts of the systematic cellular remodeling are driven by, or regulated by, NO• remains to be fully defined.

Especially, mitochondrial metabolism in activated macrophages is tightly regulated to enable immune effector functions [14,15]. Previous studies have shown that NO• (or NO-derived metabolites) can impair abundance or activity of several key mitochondrial enzymes. For example, NO• exposure reduces total levels of the electron transport chain (ETC) complex I subunits [16,17] or causes their inhibition by post-translational modification [18,19], and NO•-mediated modifications strongly inhibit α-ketoacid dehydrogenase complexes, including pyruvate dehydrogenase (PDHC), oxoglutarate dehydrogenase (OGDH), and branched-chain α-ketoacid dehydrogenase (BCKDH) [9,10,17,20,21]. These findings on individual targets led to the idea that NO• can act as a master regulator that coordinates the sequence of inhibition of mitochondrial metabolism at several steps, causing shifts in cellular redox state and dynamic changes in immunoregulatory mitochondrial metabolites in a tightly regulated manner. Given the essential role of mitochondrial metabolism, and the observations that classically activated macrophages are highly functional and highly viable despite having high RNS levels, we also hypothesized that macrophages deploy coordinated counter-regulatory mechanisms to control NO•-mediated metabolic inhibition and limit excess redox stress during classical activation.

Here we applied a multi-omic strategy to (i) define NO•-dependent regulation at a systems level and (ii) identify molecular mechanisms that balance and fine-tune the metabolic inhibition caused by NO• during classical activation. We integrated quantitative proteomic and transcriptomic profiling across primary and immortalized murine macrophages, with genetic modulation of endogenous NO• production and exogenous NO• donor treatment, to comprehensively map out NO•-mediated responses and NO•-immune signaling crosstalk. This approach revealed selective NO•-dependent remodeling of mitochondrial pathways and identified aldo-keto reductase family 1 member A1 (AKR1A1), an oxidoreductase enzyme, as a robustly NO• and stimulation co-induced response. We subsequently performed targeted biochemical assays to define the mechanism by which AKR1A1 enzymatically modulates NO•-derived inhibitory effects. Together, this study provides a systems-level characterization of NO•-dependent remodeling and demonstrates a molecular mechanism that enables macrophages to manage NO•’s effect within the context of regulated metabolic adaptation.

## RESULTS

### Global proteomic profiling reveals NO•-dependent remodeling in classically activated macrophages

To assess the regulatory impact of NO• during macrophage activation, we performed label free quantitative proteomics in primary murine bone marrow-derived macrophages (BMDMs) stimulated with LPS/IFNγ for 48 hours. To distinguish NO•-dependent effects from those arising from classical activation alone, we included conditions with or without knockout of endogenous iNOS and with or without treatment of the NO• donor, DETA-NONOate (Figure 1A).

**Figure 1.**
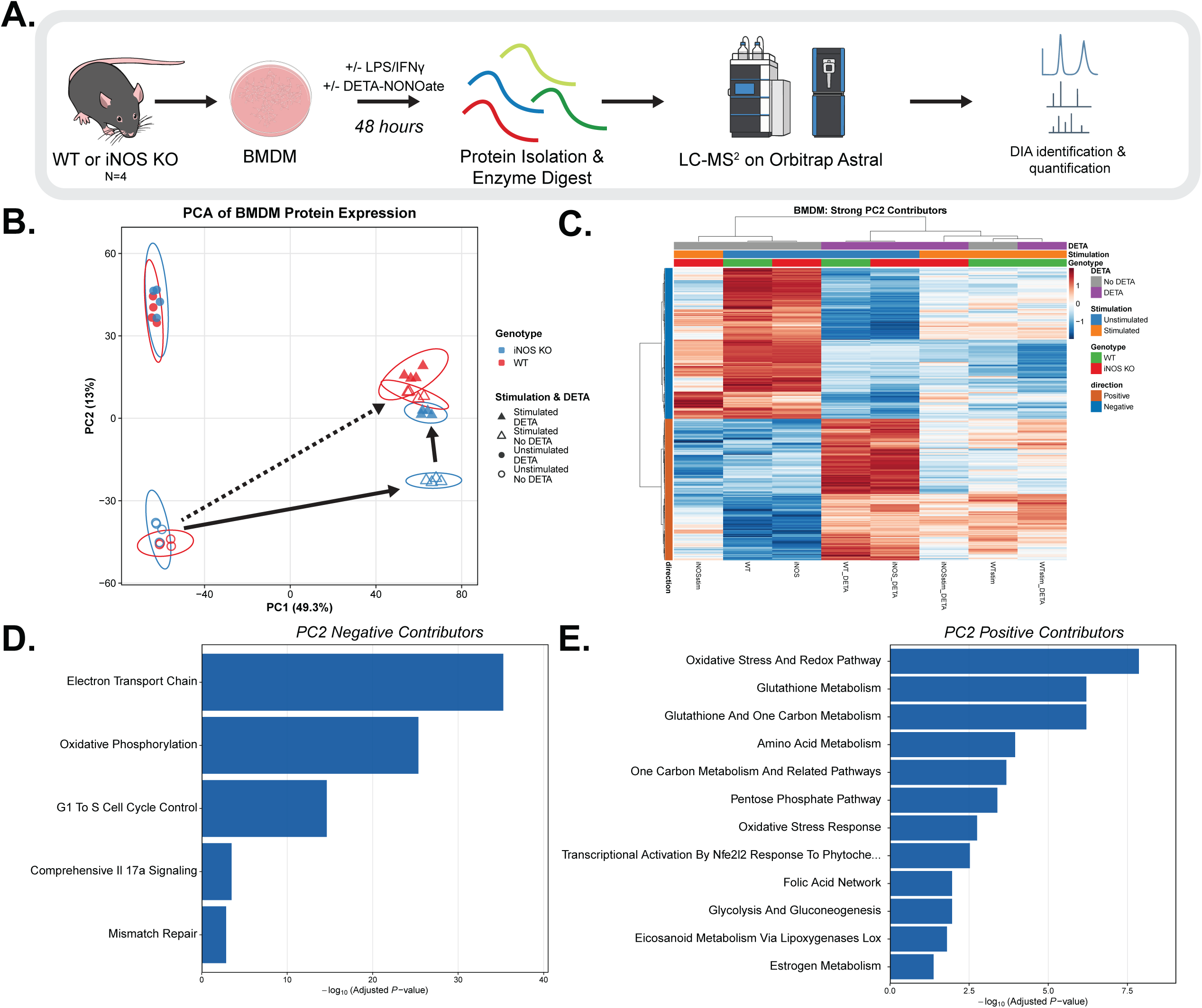
Global proteomic profiling reveals NO•-dependent remodeling in primary bone marrow-derived macrophages. **(A)** Experimental design and proteomic workflow for BMDMs from wildtype (WT) or iNOS knockout (iNOS KO) mice that were stimulated with or without LPS/IFNγ for 48-hours with or without DETA-NONOate, n = 4 biological replicates per genotype. **(B)** Principal component analysis (PCA) of all proteins. PC1 and PC2 shown with 95% confidence ellipses. **(C)** Heatmap showing Z-score of top positive or negative PC2 loading proteins across conditions (n = 432). Hierarchical clustering based on Pearson correlation distance (Ward.D2 linkage). Color scale: blue (low) and red (high) expression relative to mean. **(D-E)** Pathway enrichment analysis (Mouse WikiPathway 2024 via Enrichr) for PC2-negative (D) and PC2-positive (E) contributors.

Across all experimental groups, 7,937 proteins were analyzed. Principal component analysis (PCA) showed upon classical activation, proteome of wildtype BMDMs is substantially remodeled, with the top two principal components capture majority of the changes (Supplemental Figure 1A). At baseline, iNOS knockout (KO) cells show proteomic profile inseparable from wildtype, consistent with the absence of iNOS expression at baseline; upon stimulation, proteome shifts substantially and wildtype and iNOS KO separate significantly, moreover, such difference can be rescued by exogenous NO• donor treatment, suggesting NO• play an important role in regulating the proteomic remodeling in response to classical activation. Notably, this analysis revealed the normal proteomic remodeling upon classical activation can be decomposed into (i) NO• mediated changes along major principal component 2 (PC2), and (ii) largely NO•-independent changes induced by stimulation, which is mainly along PC1 (Figure 1B), as supported by that all four stimulated conditions, regardless of endogenous NO• production or exogenous NO• show high PC1 score whereas all four unstimulated conditions show low PC1 score (Supplemental Figure 1B). Comparing the NO• mediated shift along PC2, we observed that treatment of NO• donor causes similar but greater shift in unstimulated state than stimulated state, suggesting crosstalk between NO• and immune signaling, and that stimulation may activate some mechanisms limiting and balancing NO•’s effect. Additionally, we observed minor portion of NO•-dependent regulation along PC3, and similar cross talk between NO• and immune signaling indicated by stimulation state-dependent shift along PC3 (Supplemental Figure 1C). Overall, the result demonstrated NO• is a major regulator mediating macrophages’ response to classical activation and identified a set of NO•-specific regulatory effects. We focused subsequent analyses on this subset, which is mainly described by PC2.

To define proteins driving the PC2 separation, we examined PC2 loadings and identified 209 positive and 223 negative contributors using a >2 standard deviation threshold (Supplemental Figure 1D). Negative contributors generally show higher abundance in stimulated iNOS KO macrophages relative to wildtype and are reduced by the treatment of NO• donor, whereas positive contributors of PC2 were increased by the endogenous NO• production

(comparing stimulated wildtype cells to iNOS KO cells or unstimulated wildtype cells) or by the treatment of exogenous NO• donor (Figure 1C). This dynamic bidirectional shift indicates that NO• signaling robustly alters defined protein networks.

Enrichment analysis of these PCA-defined subsets revealed coordinated suppression of mitochondrial metabolism, particularly the electron transport chain (the top downregulated pathway by NO•) and the significant induction of antioxidant and stress-response pathways (Figure 1D and E). Proteins within these groupings included ETC complex subunits (e.g., NDUFB7, SDHB, COX7A2) and antioxidant enzymes (e.g., GPX4, PRDX1, TXNRD1; consistent with previous literature [22,23]) as well as enzymes supporting the synthesis of redox defense molecules (e.g. GCLC, SLC7A11, G6PDX), respectively [22,23]. The reciprocal enrichment of mitochondrial and antioxidant signatures suggests that metabolic remodeling and redox adaptation are core elements of the NO•-driven response.

### Cross-model proteomic analysis supports NO•-dependent remodeling in macrophage-like cells

To determine whether NO•-dependent remodeling observed in BMDMs was conserved across models, we conducted TMTpro-based quantitative proteomics in RAW264.7 macrophage-like cells stimulated with LPS/IFNγ for 48 hours, with or without iNOS KO (Figure 2A). Across all conditions, 5,237 proteins were analyzed. Hierarchical clustering (Figure 2B) revealed that out of these 5,237 proteins, a cluster of 1,522 proteins are generally induced upon stimulation in a NO•-dependent manner (iNOS KO limits the stimulation-induced upregulation), a cluster of 1,878 proteins are downregulated upon stimulation in a NO•-dependent manner, and another cluster of 998 proteins show the pattern of stimulation-induced change that is largely NO•-independent. Clustering across samples show consistent observations as in BMDMs. In unstimulated state the proteomic profile in wildtype and iNOS KO are very similar, stimulation causes substantial shifts and separation between iNOS KO and wildtype. These results confirmed that NO• production via iNOS was a major driver of proteomic reprogramming upon classical activation, supporting the idea that NO• induces systematic reprogramming rather than model-specific effects.

**Figure 2.**
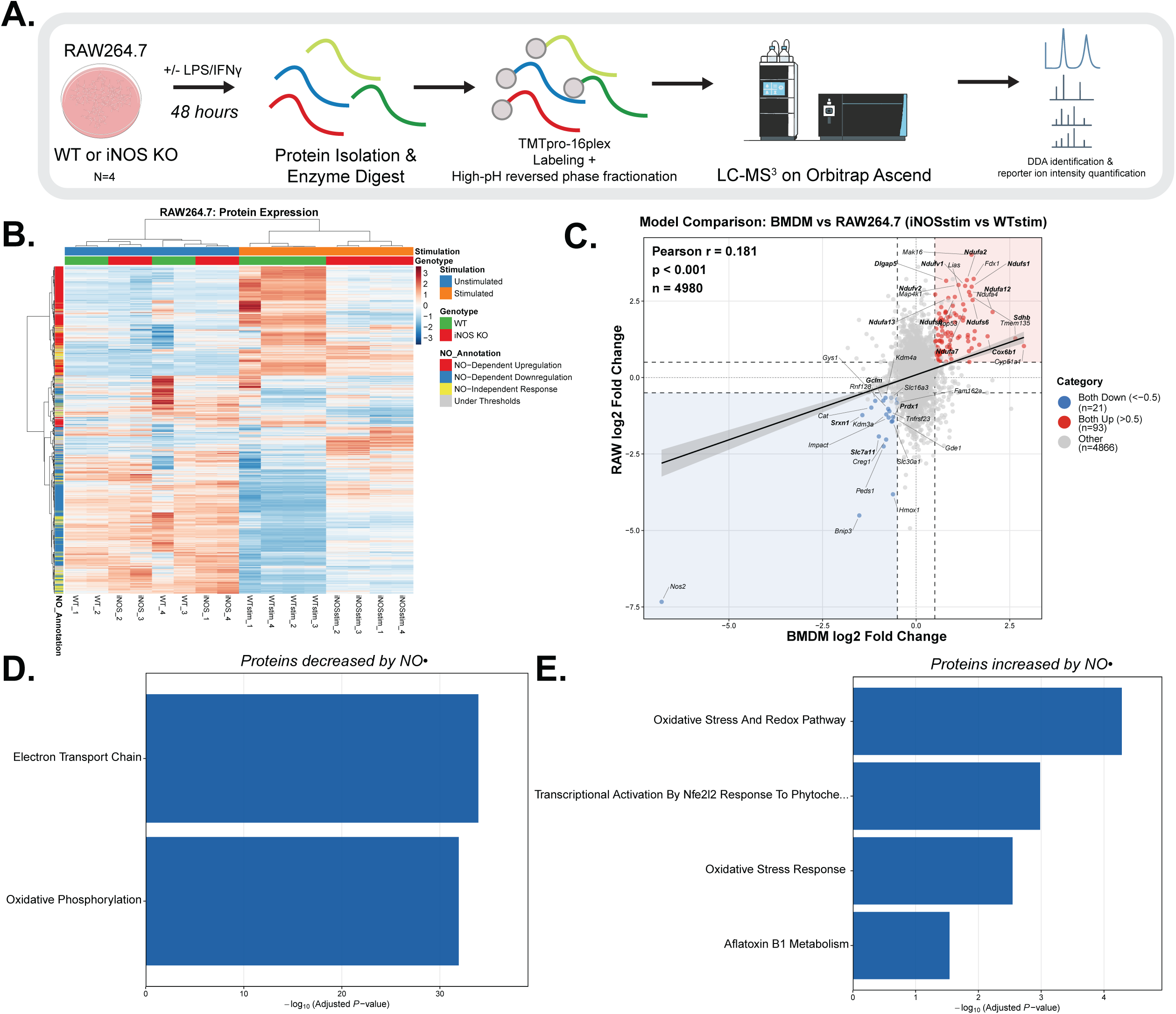
PCA-defined NO•-dependent proteomic signatures are preserved across cell models. **(A)** Experimental design and proteomic workflow for RAW264.7 macrophage-like cells. Wildtype (WT) or iNOS knockout (KO) RAW264.7 cells were stimulated with or without LPS/IFNγ for 48 hours. TMTpro 16-plex labeling with 4 biological replicates per condition. **(B)** Heatmap showing Z-score normalized expression of all proteins across condition (individual replicates designated by “_1” through “_4” suffix). Hierarchical clustering based on Pearson correlation distance (Ward.D2 linkage). Color scale: blue (low) and red (high) expression relative to mean. Rows categorized by response pattern: NO-dependent upregulation (red; WT stimulation effect > 0.4 AND NO-specific effect > 0.5); NO-dependent downregulation (blue; WT stimulation effect <-0.4 AND NO-specific effect <-0.5); NO-independent response (yellow; |WT stimulation effect| > 0.4 AND |iNOS KO stimulation effect| > 0.4 AND |NO-specific effect| < 0.5); under thresholds (grey; not meeting criteria above) **(C)** Cross-model correlation analysis comparing iNOS-dependent proteomic changes (log2 fold change: stimulated iNOS KO versus stimulated WT) between BMDM and RAW264.7 datasets for overlapping proteins. Dashed lines indicate |log2 fold change| = 0.5 threshold. Shaded regions highlight concordant quadrants. Points are colored by significance: both models significant (p-adj < 0.05) and concordantly increased (red) or decreased (blue) with |log2FC| > 0.5 in both datasets. Pearson correlation with linear regression shown (95% CI as shaded band). **(D-E)** Pathway enrichment analysis (Mouse WikiPathway 2024 via Enrichr) for concordantly decreased (D) or increased (E) proteins from panel C.

To directly compare NO•-dependent regulation between models, we analyzed log_₂_ fold changes for the stimulated iNOS KO versus corresponding wildtype stimulated conditions. Despite inherent biological differences between primary and immortalized cells [24], a modest but statistically significant correlation (r = 0.181, p-adj < 0.001) was observed (Figure 2C). In general, the effect size of iNOS is much greater in RAW264.7 cell lines, likely due to the greater NO• production and the change in proliferation status (in RAW264.7 cells, proliferation is inhibited by stimulation induced NO• production, whereas in BMDMs, cells are non-proliferating regardless of genotype or stimulation status) [8].To examine the robust NO•-mediated effects, we further filter the results by criteria that the effects are statistically significant (p-adj < 0.05) in either model and sufficient in size to be biologically meaningful (|log_2_-fold change| > 0.5). This identified 134 proteins, with the majority (114 proteins) showing consistent significant changes across the two cell models. Among the consistently decreased proteins by NO•, the top enriched process is the ETC and among the consistently increased protein, the top enriched process is oxidative stress and redox pathways (Figure 2D and E). Together, these findings reinforce that these are core NO•-dependent regulatory patterns.

### Cross-omic comparison reveals the role of transcriptional regulation in NO•-mediated remodeling

We next investigated to what extent the systematic proteomic remodeling driven by NO• is mediated through transcriptional mechanism. Bulk RNA sequencing performed in RAW264.7 cells under equivalent genotype and stimulation conditions (WT and iNOS KO +/- LPS/IFNγ) identified 3,312 differentially expressed genes (adjusted p-value < 0.05, |log_2_-fold change| > 1.0, upregulated = 1,838, downregulated = 1,474) (Figure 3A). The significant changes indicate NO• plays an important role in remodeling at the transcriptional level as well. When comparing global iNOS-dependent responses between proteomic and transcriptomic datasets, we observed a significant positive correlation (r = 0.568, p < 0.001). The highest portion of the proteins show consistent significant changes by NO• at protein and transcript level (35%), followed by those only changes significantly at the transcript level (26.1%), and only significantly at protein level (16.3%). This indicates that NO• induces coordinated and substantial changes across molecular levels (Figure 3B).

**Figure 3.**
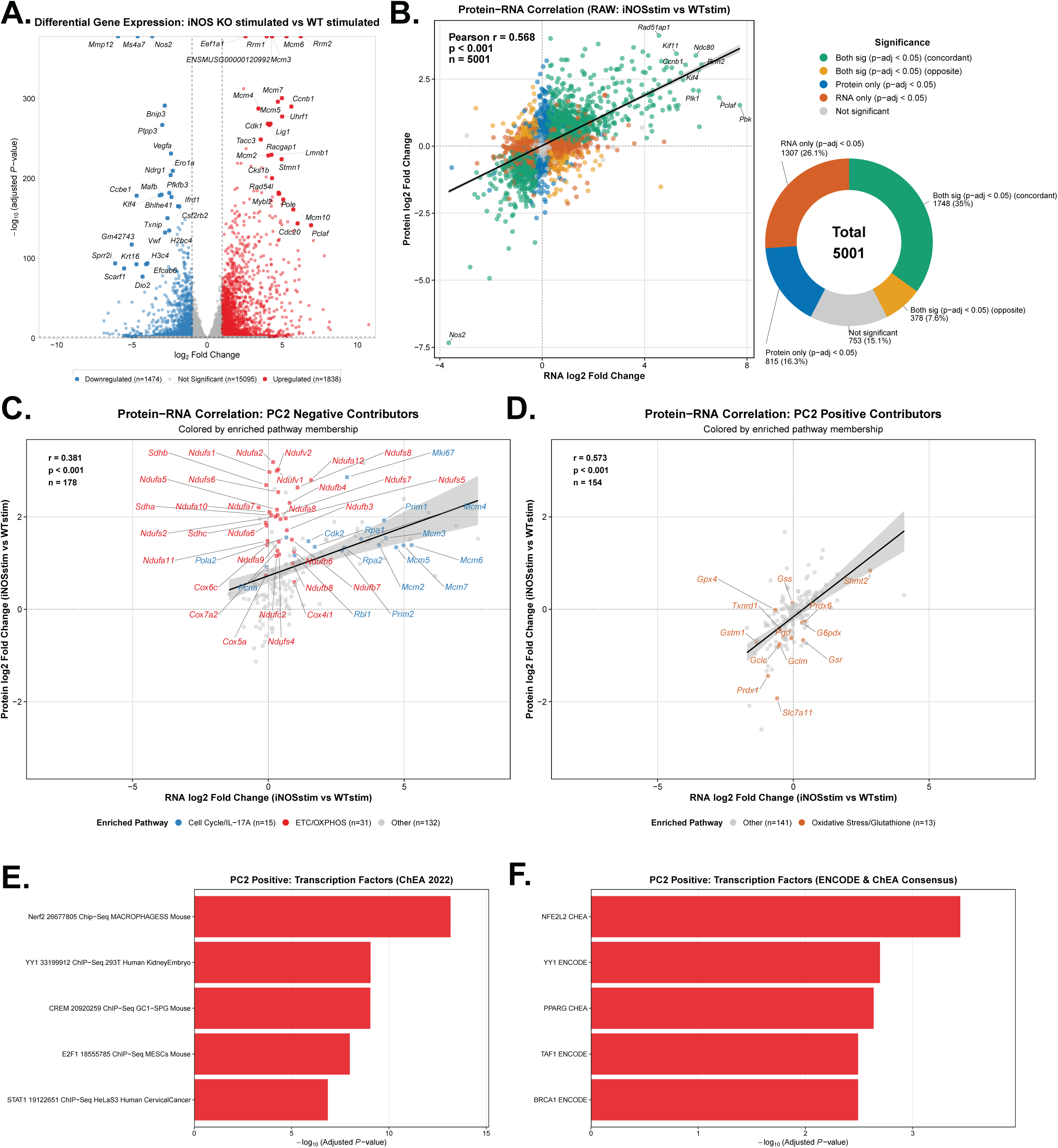
The effect of NO• in transcriptomic remodeling and its role in mediating major proteomic changes. **(A)** Volcano plot showing differentially expressed genes in stimulated iNOS knockout (iNOS KO) versus stimulated wildtype (WT) RAW264.7 cells (LPS/IFNγ, 48 hours; n = 3 biological replicates per genotype). Genes meeting significance thresholds (p-adj < 0.05 and |log2 fold change| > 1.0) are colored by direction of change: upregulated in iNOS KO (red), downregulated in iNOS KO (blue), not significant (grey). Dashed lines indicate thresholds at p-adj = 0.05 (horizontal) and |log2FC| = 1 (vertical). Top 25 genes per direction (ranked by-log10(p-adj) × |log2FC|) are labeled. **(B)** Cross-omic correlation analysis comparing iNOS-dependent proteomic and transcriptomic changes (log_2_ fold change: stimulated iNOS KO versus stimulated WT) in RAW264.7 cells for genes with corresponding protein measurements (n = 5,001). Points colored by concordance category based on statistical significance (p-adj < 0.05): protein only (blue), RNA only (orange), both with opposite direction (yellow), both concordant (green), or neither (grey). Linear regression with 95% confidence interval (shaded region). Doughnut chart shows category distribution (%) with counts in each category. Top 10 concordant genes labeled. **(C-D)** Cross-omic correlation for PCA-defined NO-regulated protein subsets. (C) PC2-negative contributors. Points colored by pathway membership based on enrichment analysis: ETC/OXPHOS (red), Cell Cycle/IL-17A signaling (blue), or other pathways (grey). All pathway-annotated genes are labeled. (D) PC2-positive contributors. Points colored by pathway membership: Oxidative Stress/Glutathione metabolism (orange) or other pathways (grey). All pathway-annotated genes are labeled. Linear regression with 95% confidence interval shown (shaded region) for C–D. **(D-E)** Transcription factor pathway enrichment analysis (E: ChEA 2022; F: ENCODE and ChEA Consensus) using Enrichr among PC2-positive contributors from D.

To examine whether processes that are most regulated by NO• are regulated through transcriptional mechanisms or other mechanisms, we focused on the top contributors to PC2 as identified in Figure 1, as these subsets represent proteins most strongly associated with NO•-dependent variance. Among the top negative contributors to PC2 (downregulated by NO•), the protein and transcript show a significant but moderate positive correlation overall (r = 0.381, p < 0.001, Figure 3C). The most enriched pathway, ETC and oxidative phosphorylation proteins, change more substantially at protein level than transcript level, suggesting non-transcriptional mechanism contributing to the NO•-driven decrease of these proteins. While the second most enriched pathway, cell cycle control and IL-17A signaling, are more significantly regulated at the transcriptional level (Figure 3C). Among the top positive contributors to PC2 (upregulated by NO•), the correlation between protein and transcript is stronger (r = 0.573; p < 0.001, Figure 3D). Furthermore, the top enriched process, oxidative stress and redox pathway, is consistently regulated at transcript and protein level, suggesting important role of transcriptional regulation. We then conducted enrichment analysis to identify possible transcriptional factors that may mediate their induction. The transcription factor, NRF2, was consistently identified as a top hit (Figure 3E and F), suggesting a likely axis mediating the NO•-dependent induction of these transcripts and proteins upon classical activation.

### Complex-specific remodeling of the electron transport chain

Having identified mitochondrial metabolism as the most enriched pathway among those negatively regulated by NO• (Figure 1D), we next examined whether NO• affected specific ETC complexes or caused broad ETC subunit suppression. Restricting correlation analyses to the ETC-annotated proteins from the KEGG Pathway confirmed strong cross-model correlation (r = 0.800, p < 0.001). Among the most strongly and consistently negatively regulated by iNOS expression are most of complex I and complex II components and many core components of complex IV (whereas its assembly factors do not follow this pattern, Supplemental Figure 2A), but not complex III or complex V (Figure 4A), Comparing across proteomics and transcriptomics in RAW264.7 cells, most components of complexes III, IV and ATPases show consistent changes at transcript and protein level, in contrast, complex II components are regulated by protein level regulation independent of transcriptional changes, while complex I components are regulated by both transcriptional and non-transcriptional processes (Figure 4B).

**Figure 4.**
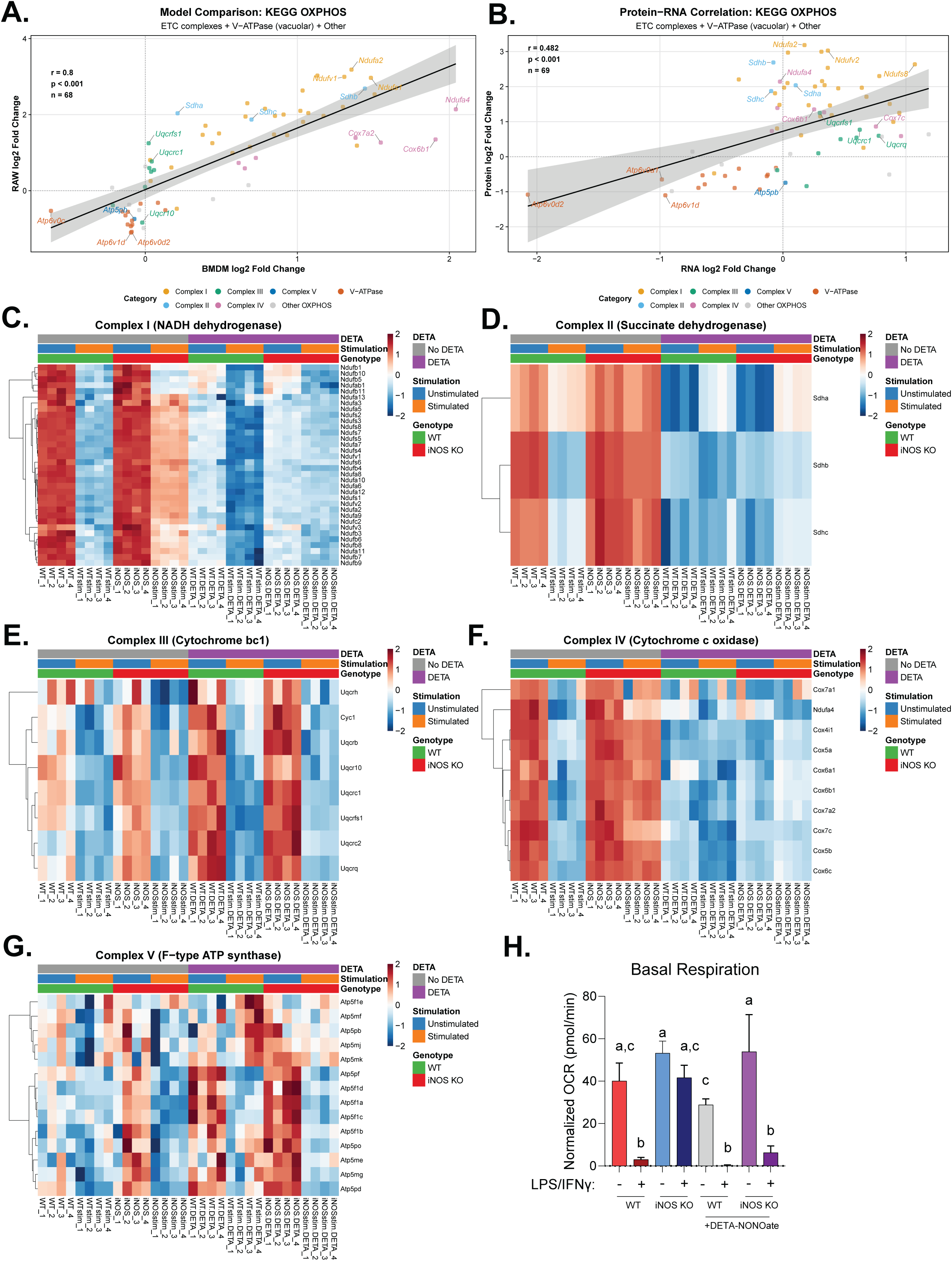
NO• drives complex-specific remodeling of the mitochondrial electron transport chain. **(A)** Cross-model proteomic correlation for ETC/OXPHOS proteins comparing iNOS-dependent changes (log_2_ fold change: stimulated iNOS knockout (KO) versus stimulated wildtype (WT)) between BMDM (x-axis) and RAW264.7 cells (y-axis). Dots represent 68 proteins defined by KEGG OXPHOS pathway that are quantified in both models, color coded by ETC Complex assignment; V-ATPase subunits (vacuolar/lysosomal) shown separately from mitochondrial Complex V (F-type ATP synthase). Linear regression with 95% confidence interval (shaded region). **(B)** Cross-omic correlation for ETC/OXPHOS proteins comparing iNOS-dependent transcriptomic (x-axis) and proteomic (y-axis) changes in RAW264.7 cells (n = 69 genes with measurements in both datasets). Points colored by ETC complex as in (A). Linear regression with 95% confidence interval shown. **(C–G)** Heatmaps showing changes in each of the ETC complexes: (C) Complex I, (D) Complex II, (E) Complex III, (F) Complex IV, (G) Complex V, across eight experimental conditions: unstimulated (unstim) or stimulated (stim) with LPS/IFNγ for 48 hours (stim) in WT or iNOS KO genotypes; +/- DETA-NONOate (DETA). Colors represent row-wise Z-score normalized protein abundance (blue = decreased, red = increased relative to row mean). Rows (proteins) clustered by Pearson correlation; columns ordered by experimental condition. n = 4 biological replicates per condition (individual replicates designated by “_1” through “_4” suffix). **(H)**. Normalized oxygen consumption rate (OCR) of BMDM across experimental conditions as described in C-G. Data represent mean + standard deviation, n = 3 biological replicates. Statistical analysis by one-way ANOVA with Tukey’s post hoc test for multiple comparisons; bars with different letters indicate statistically significant differences (p < 0.05).

While the strong shifts caused by iNOS KO in stimulated macrophages indicate NO• plays a crucial role in the remodeling of ETC upon classical activation, the activation signals (LPS/ IFNγ) can induce broad responses beyond NO• production by iNOS. To further delineate the direct effect of NO• from other effects of stimulation, we carefully compared the impacts of exogenous NO• donor treatment in unstimulated or stimulated wildtype and iNOS KO BMDMs. Interestingly, this revealed complex-specific regulatory pattern and crosstalk between NO• and LPS/IFNγ signaling. The downregulation of complex I show additive effect of NO• and stimulation: comparing to the corresponding unstimulated condition (regardless of whether or not treated with NO• donor or iNOS KO), stimulation causes decrease across the complex I subunits; on the other hand, comparing to the corresponding untreated conditions, the four NO• donor treated conditions all show additional decrease across complex I subunits (Figure 4C). This suggests NO•-dependent and NO•-independent stimulation-induced mechanisms both contribute to the downregulation of complex I upon activation. In contrast, complex II and some complex IV components show primarily NO• dependent downregulation, as NO• donor treatment is sufficient to cause similar decrease in these proteins as stimulation in wildtype, and in conditions where endogenous iNOS is knocked out, stimulation does not cause consistent decrease compared to its corresponding unstimulated condition (Figure 4D and F). Complex III subunits mainly show minor stimulation-dependent, NO•-independent downregulation in BMDM, as endogenous NO• production or exogenous NO• has no major additional effects (Figure 4E). Finally, most complex V subunits do not show consistent decrease by stimulation or NO•, some are even slightly increased (Figure 4G).

To examine overall mitochondrial changes, we used an imaging approach. Despite the significant decrease in many ETC components, stimulation was not found to cause significant changes in mean mitochondrial volume, mitochondrial sphericity, or total mitochondria number (Supplemental Figure 3A-D). This result, consistent with the observation that the downregulation is specific to certain complexes, further indicates the regulation is at the individual ETC complex level rather than general decrease of mitochondria.

To examine the main function of ETC, cellular respiration, we measured oxygen consumption rates. Consistent with previous reports [16,17], classical activation causes profound decrease in basal respiration rate. Intriguingly, we found this respiration inhibition depends on both NO• and stimulation; neither NO• donor treatment alone nor stimulation alone of iNOS knockout cells is sufficient to cause the respiration inhibition (Figure 4H). Given the additive effect of NO• and stimulation on complex I components, as well as the specific effect of NO• on complex II and complex IV and the specific effect of stimulation in complex III, it is likely that it takes two hits to inhibit ETC to an extent that becomes rate limiting. Thus, NO• is required, but not sufficient, in the strong inhibition of mitochondrial respiration associated with macrophage classical activation.

### Upregulation of Akr1a1 is induced by stimulation in a NO-dependent manner

We next sought to investigate proteins that are upregulated by NO•. Enrichment analysis showed these are enriched for oxidative stress and redox pathways (Figure 1E and 2E), suggesting these responses could play an important role in balancing some NO•-mediated effects in redox or free radical stress providing a metabolic adaptation. Among the proteins that most consistently upregulated upon stimulation in a NO•-dependent manner (as indicated by significant decrease by iNOS knockout across the omic dataset), aldo-keto reductase family 1 member A1 (AKR1A1) emerged as a top hit (Figure 5A). Previous studies have identified AKR1A1 has reductase activities towards S-nitroso-glutathione and S-nitroso-Coenzyme A (SNO-CoA; a reductive intermediate formed by nitrosylation of coenzyme A) [25,26]. Given its activities in metabolizing these important low molecular weight thiol containing molecules, which play a crucial role in mediating NO•-induced proteins modifications including s-nitrosylation of cysteine residues [1] and other more recently identified regulatory modifications [20,21], we hypothesized that the NO•-dependent upregulation of AKR1A1 can provide a check-and-balance mechanism to tightly regulate the remodeling upon macrophage classical activation.

**Figure 5.**
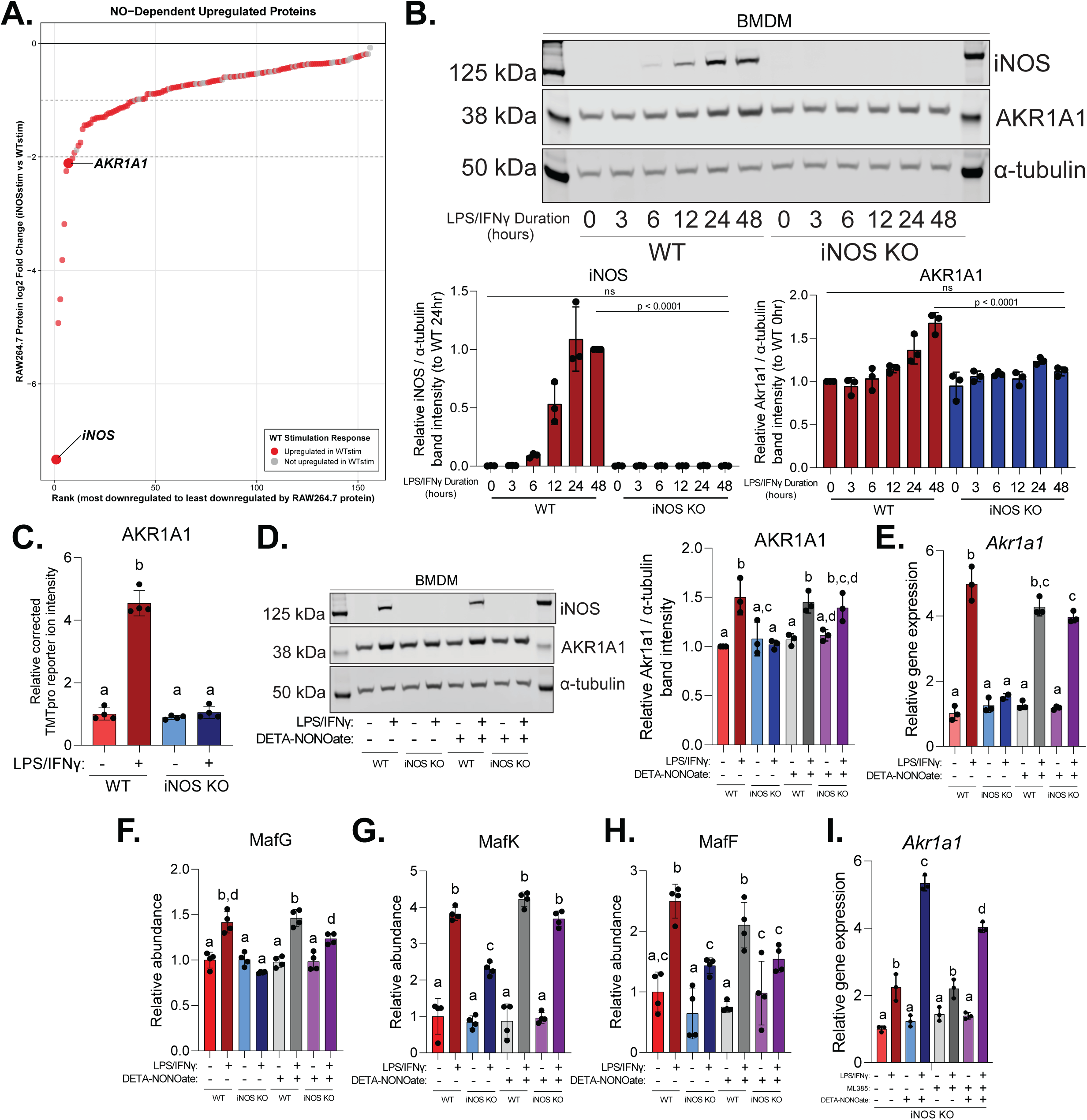
***Akr1a1* induction requires both classical activation and NO• signaling and is partially NRF2-dependent (A)** Proteins significantly (p-adj < 0.05) decreased in iNOS KO vs WT (both LPS/IFNγ-stimulated) across all three datasets (BMDM proteomics, RAW264.7 proteomics, RAW264.7 RNA-seq), ranked by RAW264.7 protein fold change. Red highlights proteins significantly induced by LPS/IFNγ in WT cells (p-adj < 0.05, log2FC > 0), indicating NO•-inducible proteins. **(B)** Top: Representative immunoblot showing AKR1A1 and iNOS protein abundance in WT and iNOS KO BMDMs stimulated with LPS/IFNγ for 0, 6, 12, 24, or 48 hours. Bottom: Quantification of AKR1A1 and iNOS protein abundance (normalized to α-tubulin). Data represents mean +/-standard deviation (SD) from n = 3 independent experiments. **(C)** Relative abundance of AKR1A1 protein from RAW264.7 cell proteomics dataset. **(D)** Left: Representative immunoblot showing AKR1A1 and iNOS protein abundance in WT and iNOS KO BMDMs under four conditions: unstimulated and stimulated with 48-hour LPS/IFNγ alone, 48-hour 200 μM DETA-NONOate alone, or 48-hour LPS/IFNγ and DETA-NONOate. Right: Quantification of AKR1A1 protein abundance (normalized to α-tubulin), confirming proteomic measurement. Data represents mean +/- SD, n = 3 independent experiments. **(E)** Relative mRNA expression of *Akr1a1* measured by RT-PCR from cells treated as in D. Expression normalized to *Hnrpab* reference gene using ΔΔCt method. Data represents mean +/- SD, n = 3 biological replicates. **(F-H)** Relative abundance of MafG (F), MafK (G), and MafF (H) from BMDM proteomics dataset. **(I)** Relative mRNA expression of *Akr1a1* in iNOS KO BMDMs stimulated with LPS/IFNγ +/-DETA-NONOate +/- ML385 (Nrf2 inhibitor, 10 μM) for 48 hours. Expression normalized to *Hnrpab*. Data represents mean +/- SD, n = 3 biological replicates. <u>Statistics</u>: For panels B-J, statistical comparisons were performed using one-way ANOVA with Tukey’s post hoc test for multiple comparisons. Bars with different letters indicate statistically significant differences (p < 0.05). For panel B, ns indicates not significant.

To validate the NO•-dependent upregulation of AKR1A1 and further define its dynamics, we measured the changes in AKR1A1 protein level over a time course of LPS/IFNγ stimulation in wildtype or iNOS KO BMDMs. AKR1A1 protein abundance increased in wildtype cells but remained unchanged in iNOS KO cells (Figure 5B). The temporal changes are in strong correlation with iNOS induction and extracellular nitrite accumulation (Figure 5B and Supplemental Figure 4A). This indicates that endogenous NO• production is required for AKR1A1 induction during classical activation. Similarly, the compete dependence of ARK1A1 upregulation on iNOS was consistently observed in RAW264.7cells (Figure 5C).

To determine whether NO• alone was sufficient for AKR1A1 induction, wildtype or iNOS KO macrophages were exposed to the NO• donor, DETA-NONOate. We that NO• donor treatment did not cause significant increase of AKR1A1, demonstrating that NO• without classical activation signal is insufficient to cause the induction. In contrast, providing NO• during classical activation, either through endogenous iNOS activity or exogenous donor treatment, resulted in robust AKR1A1 induction (Figure 5D). RT-PCR analysis of the transcript level showed a pattern very similar to protein level that the strong induction of *Akr1a1* requires both stimulation signal and the presence of NO• (Figure 5E). These findings indicate that the upregulation of Akr1a1 upon classical activation is mainly through a transcriptional mechanism, where NO• is strictly required but not sufficient.

To further delineate signal integration, iNOS KO macrophages were treated with different combinations of LPS, IFNγ, and DETA-NONOate. Interestingly, strong *Akr1a1* induction required concurrent presence of all three components. The combination of LPS and NO• causes a partial induction. IFNγ alone, or in combination with NO•, does not cause a significant induction of *Akr1a1*, but it synergizes with LPS and NO• to cause a strong (∼5x) *Akr1a1* upregulation (Supplemental Figure 4B). This suggests a NO-independent effector downstream of LPS signaling, and amplified by IFNγ, is required for the transcriptional induction of *Akr1a1*.

Our enrichment analysis revealed that the proteins induced upon classical activation in a NO•-dependent manner are enriched for targets of transcriptional activation by NRF2 pathway, which is consistent with previous literature [23]. Therefore, we hypothesize that NRF2 contributes to the transcriptional upregulation of *Akr1a1*. Supporting this hypothesis, we found known NRF2 binding partners, small MAFs (MafG, MafK, and MafF) [27], are increased upon classical activation in a way that involves the contribution of both NO• and LPS/IFNγ stimulation (Figure 5F-H). To test this hypothesis, we treated cells with a NRF2 inhibitor, ML385. ML385 treatment does not cause reduction of *Akr1a1* at baseline, but it caused a significant reduction in the stimulation-induced upregulation of *Akr1a1* in the presence of NO (Figure 5I), similarly, a significant reduction was observed for the induction of *Hmox1*, a canonical NRF2 target (Supplemental Figure 4C), supporting that NRF2 contributes to the transcriptional control upon macrophage activation.

Together, these regulatory requirements and mechanism position *Akr1a1* at the convergence of toll-like receptor and cytokine signaling, and oxidative stress responses, as schematically represented in Supplemental Figure 4D. These findings establish the regulatory context for investigation of AKR1A1’s biochemical role in controlling NO•-mediated metabolic inhibition.

### Akr1a1 counteracts NO-mediated inhibition of PDHC by enzymatic reduction of S-nitroso-CoA

Previous studies have demonstrated a major function of AKR1A1 as an SNO-CoA reductase [25,26,28], and it has significant biological impacts in other cell types, such as in renal proximal tubules in the setting of acute kidney injury [29], in myocardium in the setting of myocardial infarction [30], and in hepatocytes with a role in cholesterol regulation [31]. In macrophages, SNO-CoA mediates the strong inhibition of mitochondrial alpha-ketoacid dehydrogenases upon classical activation, by specifically delivering NO•-driven covalent modifications onto the catalytic lipoic arms of their CoA-binding E2 subunits, as shown by our recent work [20,21]. Such inhibition drives the dynamic remodeling of the tricarboxylic acid (TCA) cycle, which modulates the levels of immunoregulatory metabolites and significantly orchestrates macrophages functions during immune responses [10,17]. Therefore, here we focus on one of the most important mitochondrial alpha-ketoacid dehydrogenases, pyruvate dehydrogenase complex (PDHC), which is the gatekeeping enzyme of complete glucose oxidation in the mitochondria. We hypothesize that by reducing SNO-CoA, AKR1A1 can control the NO•-driven inhibition of PDHC, by reducing the modifications of its lipoic arm (and thus better retaining its functional lipoic arm) (Figure 6A).

**Figure 6.**
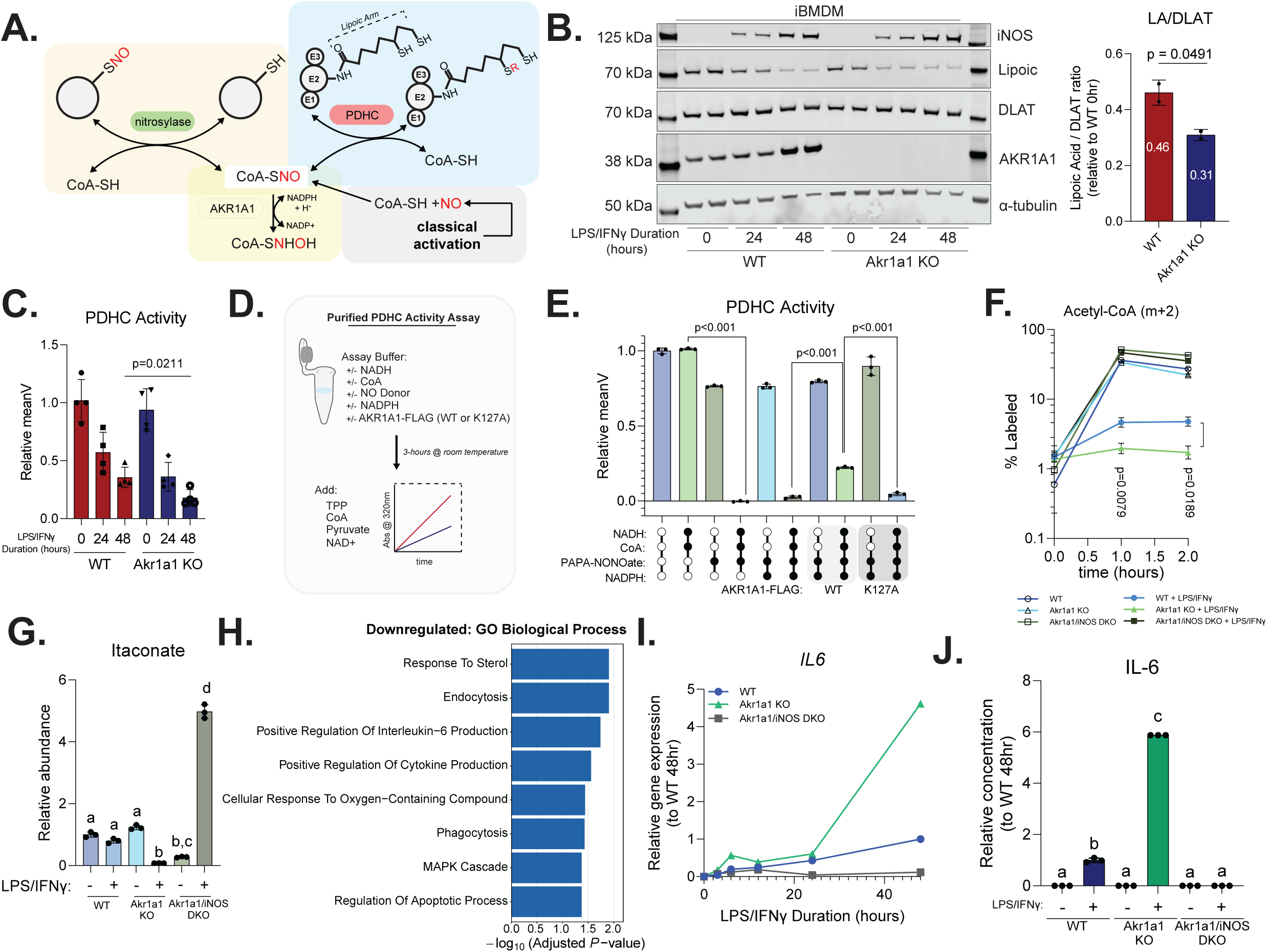
Akr1a1 regulates metabolic and functional response to classical activation by counteracting NO•. **(A)** Schematic showing AKR1A1’s main enzyme activity as SNO-CoA reductase (yellow) and its mechanistic connection to NO• driven inhibition of pyruvate dehydrogenase complex (PDHC) (blue). SNO-CoA is the key molecule delivering NO-derived modifications onto the lipoic cofactor at the catalytic center of PDHC’s E2 subunits **(B)** Left: Representative immunoblot showing iNOS, DLAT (E2 subunit of PDHC), AKR1A1 protein abundance, along with functional lipoic acid detection (anti-lipoic acid antibody) at the molecular weight of DLAT in immortalized bone marrow-derived macrophages (iBMDMs). Wildtype (WT) or *Akr1a1* KO (KO) iBMDMs were stimulated with LPS/IFNγ for 0, 24, or 48 hours. Each lane represents an independent iBMDM clone (generated using independent guide RNAs for *Akr1a1* KO). Right: Quantification of functional lipoic acid to DLAT protein ratio at 24-hour timepoint, normalized to unstimulated WT condition. Data represents mean +/- standard deviation (SD), n = 2 independent clones per genotype. **(C)** Relative PDHC enzymatic activity measured in cell lysates from iBMDMs treated as in (B) at 0-, 24-, and 48-hour timepoints. Activity normalized to unstimulated WT. Data represents mean +/- SD, n = 4 measurements per genotype per timepoint (2 independent iBMDM clones assayed in 2 separate experiments)**. (D)** Experimental design schematic for in vitro PDHC–Akr1a1 co-incubation activity assays. Purified porcine PDHC was incubated for 3 hours at room temperature with indicated combinations of PDHC substrates (NADH and CoA) and PAPA-NONOate (NO• donor) +/- purified Akr1a1-FLAG (WT or K127A) +/- NADPH (Akr1a1 cofactor), followed by measurement of PDHC activity. **(E)** Relative PDHC enzymatic activity from in vitro co-incubation assays as described in (D). All components added simultaneously and incubated for 3 hours at room temperature before saturating levels of substrate and activity measurement. Activity normalized to protein only control. Data represents mean +/- SD, n = 3 independent reactions. **(F)** Percent 2-labeled acetyl-CoA from kinetic U-^13^C-D-glucose tracing in WT, *Akr1a1* KO, or *Akr1a1/iNOS* DKO RAW264.7 cells unstimulated or stimulated with LPS/IFNγ for 48 hours. Data represents mean +/- SD, n = 3 biological replicates. **(G)** Relative total abundance of itaconate from conditions described in (F). **(H)** Pathway enrichment analysis (GO Biological Process) using Enrichr among downregulated differential expressed genes from Supplemental Figure 6A. **(I)** Relative mRNA expression of *IL6* from of WT, *Akr1a1* KO, and *Akr1a1/iNOS* DKO RAW264.7 cells stimulated for 0-, 3-, 6-, 12, 24-, and 48-hours. Expression normalized to *Hnrpab*. **(J)** IL-6 measured via ELISA from media of WT, *Akr1a1* KO, and *Akr1a1/iNOS* DKO RAW264.7 cells unstimulated or stimulated for 48-hours. Data represents mean +/- SD, n = 3 biological replicates. <u>Statistics</u>: For B and C panels, statistical comparisons with unpaired two-tailed Student’s t-test. For panel E-G and J, statistical comparisons were performed using one-way ANOVA with Tukey’s post hoc test for multiple comparisons. Bars with different letters indicate statistically significant differences (p < 0.05), or exact p-value reported.

Indeed, in immortalized bone marrow-derived macrophages (iBMDMs), we found *Akr1a1* KO significantly slowed down the gradual loss of functional (*unmodified*) lipoic arm on PDHC E2 subunit (Dihydrolipoamide S-acetyltransferase, DLAT), which is induced by LPS/IFNγ stimulation (Figure 6B). Consistently, at the enzymatic activity level, the inhibition of PDHC was significantly accelerated and exaggerated in *Akr1a1* KO cells upon classical activation (Figure 6C). These indicate that induction of *Akr1a1* can be protective for PDHC activity by tempering its inhibition over the time course to immune response.

To mechanistically test whether AKR1A1 modulates PDHC inhibition via SNO-CoA reduction, we performed in vitro assays using purified PDHC co-incubated with either wildtype FLAG tagged AKR1A1 or a catalytically inactive K127A mutant [26,28] (Figure 6D and Supplemental Figure 5A). PDHC enzymatic activity was strongly inhibited by NO• donor, but only when its substrates CoA and NADH were also present during the co-incubation to generate SNO-CoA and to reduce lipoic cofactor to its reactive thiol containing form, respectively, consistent with the previously demonstrated mechanism that PDHC inhibition is specifically dependent on SNO-CoA modifying the lipoic arm [20]. Under these inhibitory conditions, wildtype AKR1A1 provides significantly, partial protection for PDHC activity, when its substrate NADPH is available, whereas the catalytically dead K127A showed no effect (Figure 6E).

To assess temporal requirements, PDHC was first exposed to PAPA-NONOate, CoA, and NADH prior to addition of AKR1A1. When AKR1A1 was added after the establishment of SNO-COA-mediated inhibition, no rescue was observed. However, when AKR1A1 was present during the initial incubation, PDHC activity was partially rescued (Supplemental Figure 5B). These results suggest that AKR1A1 acts preventively by intercepting inhibitory intermediates, but it cannot reverse the inhibition by removing the inhibitory modifications.

In cells, PDHC controls the influx from glucose into TCA cycle, and its inhibition has been shown to drive the shutdown of mitochondrial glucose oxidation at later timepoints upon macrophage classical activation [10]. To test the impact of *Akr1a1* in regulating TCA cycle flux, we generated *Akr1a1* KO and *Akr1a1*/*iNOS* double knockout (DKO) RAW264.7 cells (Supplemental Figure 5C). Tracing with U-^13^C-D-glucose showed that *Akr1a1* KO significantly exaggerated the inhibition of glucose entering TCA cycle upon stimulation, as indicated by the further loss of 2-labeling in acetyl-CoA (Figure 6F) or metabolites downstream of acetyl-CoA (e.g., cis-Aconitate) (Supplemental Figure 5D). The decrease in labeling is completely prevented by *Akr1a1*/*iNOS* DKO, indicating the stimulation-induced inhibition is dependent on NO•, and *Akr1a1* provides protection through countering the inhibitory effect of NO•.

Collectively, these findings demonstrate that AKR1A1 enzymatically reduces SNO-CoA to protect the lipoic arm of PDHC from NO•-induced modification, thereby delaying suppression of PDHC activity and the inhibition of TCA flux upon classical activation. Of note, while we focused on PDHC, other lipoate-dependent enzymes (e.g., OGDH and BCKDH) [20,21] are likely similarly protected by AKR1A1, as they also functionally act as SNO-CoA-assisted nitrosylases [32] to self-modify in a SNO-CoA dependent manner.

### Akr1a1 regulates NO-mediated metabolic and functional remodeling in classically activated macrophages

The dynamic changes in TCA cycle during the immune response to classical activation, particularly the inhibition of PDHC, which we have shown to be a key regulatory event that drives the metabolic transition from more inflammatory early response stage to the late stage [10], is important for the tight and timely regulation of macrophage functions. One important mechanism is related to the rise-and-fall of immunoregulatory metabolite itaconate, which has been shown to regulate the dynamic changes in macrophage functions over the immune response time course via its impact on several signaling pathways [33–37]. Itaconate first accumulates at early timepoints due to the rapid induction of immune-responsive gene 1 (IRG1), then falls back to at or below baseline levels at later timepoints primarily due to NO-driven inhibition of PDHC, which shuts off influx for itaconate production from upstream [10]. Consistently, we found that *Akr1a1* KO greatly exacerbated the decrease of itaconate at 48-hours post-stimulation. To the contrary, in *Akr1a1/iNOS* DKO, where PDHC activity is not inhibited, itaconate accumulates greatly (Figure 6G). This demonstrates AKR1A1’s significant role in regulating the level of immunoregulatory metabolite during immune response by counteracting NO•.

To understand the broader impact of *Akr1a1*, we performed bulk RNA-seq analysis of wildtype and *Akr1a1* KO RAW264.7 cells (Supplemental Figure 6A). Among the top significantly enriched processes that are impacted by *Akr1a1* KO in stimulated state are cytokine production, endocytosis, and phagocytosis (Figure 6H), which are some of the major functions in activated macrophages. For targeted validation, we focused on IL-6, which is an important inflammatory cytokine produced by macrophages and has been shown to be negatively regulated by itaconate [38]. We found that in *Akr1a1* KO cells, *IL6* expression continues to increase greatly especially at later time points post-stimulation, reaching over 4-fold higher than wildtype by 48 hours. In contrast, in the *Akr1a1/iNOS* DKO cells, IL6 expression starts to fall earlier, and is much lower than wildtype by 48 hours (Figure 6I). Consistently with gene expression, IL-6 secretion in *Akr1a1* KO cells increases to over 5-fold higher than wildtype at 48 hours post-stimulation, whereas in *Akr1a1/iNOS* DKO cells, IL-6 secretion is much lower than wildtype (Figure 6J). These results support that *Akr1a1* can influence macrophage functions through counteracting NO•.

## DISCUSSION

Here using integrated multi-omic profiling and targeted analyses, we demonstrated that NO• mediates structured cellular remodeling during macrophage classical activation. Among the NO•-dependent regulations, mitochondrial metabolism was found to be the most negatively regulated process, and proteins involved in redox defense, including those conventionally categorized as “anti-oxidative stress” proteins and less recognized reductase AKR1A1, are the most positively regulated. Overall, this provides a comprehensive, quantitative characterization of NO•’s regulatory effect across transcript and protein levels, and highlighted NO• as a key master regulator coordinating many structured changes, especially those involved in redox metabolism, in the context of macrophage immune response. This is consistent with recent work by our group and others that elucidated NO•’s essential role in regulating specific pathways that are substantially remodeled during the immune response, including TCA cycle and nucleotide metabolism [8–10,17,20,21]. Importantly, both this work and previous studies support NO•’s role is truly regulatory, targeting selected processes, and is tightly regulated. This is essential for macrophages to stay highly viable and preform many crucial immune functions in a regulated manner in a cellular state where NO• production is high.

The comprehensive analysis revealed interesting regulatory logics that ensure tight control of NO•’s regulatory effects. First, many important NO•-mediated responses also require other NO•-independent signals downstream of LPS/IFNγ stimulation, demonstrating an “*and gate*” in regulation. This mode is exemplified by the strong inhibition of mitochondrial respiration and the upregulation of genes such as *Akr1a1*. Such “*and gate*” can allow cells intergrade information about immunological situation (e.g., sensing pathogen-associated molecular pattern) and metabolic status (e.g., redox state and NO• production, which is influenced by nutrient availability and microenvironment) in their response. The “*and gate*” could also enable context-specific response to NO and prevent potentially harmful non-specific effects of NO•, for instance, preventing broad inhibition of cellular respiration in cells not responding to immune stimulation in a high NO• microenvironment. Mechanistically, this regulation mode may be in part achieved by transcriptional regulators that both respond to immune stimulation and are regulated by NO•-mediated modifications or redox signaling. Results on the regulation of *Akr1a1* suggest NRF2 pathway can be a contributor, other candidate transcription factors that can mediate the requirement for concurrent LPS/IFNγ signaling and NO• include NF-kB, AP-1 and STAT1 [39]. Future in-depth studies examining them in a targeted manner will elucidate the contributions of these pathways. Another mechanism may achieve this functional “*and-gate*” is additive effects of NO• and other stimulation-induced response (as exemplified here by the downregulation of complex I), in combination with a high threshold for functional impact. Functionally for cellular respiration, there is likely to be a high threshold such that only when complex I decrease is profound enough, it becomes rate limiting for overall mitochondrial oxidation and drives its strong inhibition. This can provide a “*two-hit*” modeling in the regulation of respiration.

The second regulatory mode that highlighted by the results here is negative regulation, which provide a check-and-balance to fine tune the extent and timing of NO•’s regulatory effects, and protect against non-specific redox damage. The NO•-dependent upregulation of reductases and enzymes in redox defense demonstrated that NO• acts not only as an inhibitory signal but also as a cue to engage adaptive redox responses. Among these NO•-induced proteins, AKR1A1 emerged as one of the most consistent responses, and was then biochemically validated to temper NO•-driven inhibition of PDHC upon stimulation, through its enzyme activity to reduce SNO-CoA, thereby limiting the modification of the DLAT’s lipoic arm. AKR1A1 is likely to have similar effects in controlling other NO•-driven post-translational modifications on boarder protein targets, including the inhibitory lipoic modifications on other alpha-ketoacid dehydrogenases [20,21], and other cysteine nitrosylation (the protective effect against cysteine nitrosylation has been shown on other biological systems, but needs to be fully elucidated in the context of macrophage activation) [1,29–31], especially if they involve SNO-CoA assisted modification mechanisms. We found that the upregulation of AKR1A1, which peaks at 24-48-hours post stimulation concurrent with maximal NO• production (Figure 5B and Supplemental Figure 4A), does not fully prevent or reverse NO•-driven inhibition of PDHC, but significantly delays it. The inhibition of PDHC by NO• is not a “side effect” but rather plays a necessary regulatory role driving the metabolic transition from early to late response stage [10]. In macrophage immune response, the proper extent and the timing of responses is crucial. Often early in the response, supporting anti-pathogen defense functions is the primary concern, while later in the response, controlling inflammation and supporting functions promote healing become more important [40,41]. Either insufficient or excessive response can be detrimental [42], so precise control in regulation is critical. The upregulation of AKR1A1 and other similar proteins can provide the critical fine-tuning and ensure appropriate timing of the sequential metabolic and functional transitions. Supporting this idea, we found that knocking out *Akr1a1* significantly alters the level of immunoregulatory metabolite, itaconate, and influences *IL6* gene expression and IL-6 production. Building on this, future study will elucidate the exact mechanisms mediating AKR1A1’s impacts on inflammatory response, examine the connections of its role in fine tuning dynamic metabolic remodeling and functional remodeling, and explore the relevance and therapeutic potential of AKR1A1 (and other NO-induced reductases) in inflammatory conditions.

This study also advanced our understanding of NO•-driven regulation of mitochondrial respiration. Previous reports have demonstrated the inhibition of specific ETC components by NO• [16,17,43–45]. Surprisingly, through analyzing a comprehensive panel of perturbations of endogenous NO• production or exogenous NO• donor, we found NO• is required but not sufficient in the ETC downregulation. This work also generated a comprehensive multi-omic characterization of ETC remodeling, which demonstrated the remodeling is complex-specific. Most of complex I, II and complex IV are consistently downregulated by stimulation and NO•, whereas complex III and V are not affected or slightly increased. The changes in relative complex abundance can alter the capacity of mitochondria to oxidize different substrates and the partitioning of ETC in generating energy or reactive oxygen species (ROS). This can have important consequences in regulating macrophage functions during immune response, especially given the crucial signaling role of ROS [46]. For instance, previous works showed that during classical activation, macrophages increase ROS production by reversed electron transport chain, which requires altered activity of complex II and complex I, but does not involve direct participation of complex V [47]. Changes in compositions within the complex can also have significant regulatory effects. An inspiring recent work showed that the decrease of NDUFA4 within complex IV can amplify interferon program through mtDNA release [48]. The biological impacts of the NO•-dependent remodeling of the composition of ETC in macrophage response merit further investigation. Another intriguing question for continued investigation is the mechanisms driving the complex-specific regulation. Our data here showed the downregulation of certain complex IV subunits is mainly transcriptional, as the protein changes are highly proportional to transcriptional changes, in contrast, complex II subunits decrease at protein level without notable transcriptional changes, suggesting post translational mechanisms, while complex I shows a combined contribution of transcriptional and non-transcriptional mechanisms. NO•- and inflammation-responsive transcription factors or coactivators can mediate coordinated regulation at the transcriptional level [49], and likely candidates include PGC-1α, ERRα [50–52], or NRF1/NRF2 [7], in addition to metabolic remodeling drivers such as HIF-1α [4]. Whereas NO•-mediated disruption of iron-sulfur cluster is likely a mechanism causing the strong non-transcriptional decrease of protein abundance, which was mainly observed for complex II and to a less degree, complex I. Both complexes contain iron-sulfur clusters, and previous literature has shown that their disruption by reactive oxygen and nitrogen species causes destabilization of the proteins and ubiquitin-mediated proteasomal degradation [46,53]. In addition to the changes in total protein abundance, complex I and II have been found to be S-nitrosylated by NO• or NO•-derived compounds [18,19], which can add to the functional inhibition, and may be related to the stability change.

This work applied multi-omic approach to comprehensively characterize NO•-dependent regulation in macrophages. Such integrated systems-level approach revealed structured regulatory modules that would not emerge from target-based investigation alone and provides a useful resource and map for future discoveries. However, this study has several limitations, and points to a few future directions where follow-up study is needed. First, the regulatory effects of NO• in macrophage response across different models and biological contexts require to be further examined, as the responses to NO• may be heterogeneous depending on other environment or cellular conditions. Here to capture robust responses to NO•, we compared the proteomic changes in BMDMs and a macrophage-like cell line. The proteomic methods differ slightly between models (label free quantification for BMDMs versus TMTpro for RAW264.7 cells due to sample numbers). Nonetheless, the most highly enriched NO•-dependent signatures were largely consistent. While correlations were modest at times, the directional consistency in the proteins that are substantially affected by NO• suggests robust, structured regulation, and the variable effects likely reflect biological heterogeneity between primary and immortalized cells [24,54]. However, all this-omic analysis and later targeted investigations are in cell models, which may not fully recapitulate physiological conditions or complexity of NO• signaling. Future in vivo studies, for example, using macrophage-specific *Akr1a1* knockout mice or *in vivo* inflammatory models will be valuable to validate and extend the findings here. Second, here we systematically profiled the regulatory effects of NO• at transcriptomic level and proteomic level, while the involvement of post-translational modifications (PTMs) was only investigated in a targeted manner. Protein PTM, such as cysteine and tyrosine nitrosylation [55,56], and other post transcriptional regulation [7,48,57] are crucial layers in NO•-driven regulation. Future work to better integrate chemoproteomics and redox proteomics [57,58] capturing PTM with parallel datasets at transcript and whole protein level in the same biological context would provide a more complete picture of NO-dependent regulation. Finally, within the scope of this work, we were only able to perform in-depth investigation of some selected pathway or protein emerged from the omic analysis. The multi-omic dataset provided here can enable identification of additional regulatory axis and adaptive responses. The associations between proteome and transcriptome suggested additional regulatory mechanisms that merit follow-up and the significance of other NO•-induced redox defense responses revealed here (besides AKR1A1) merit further elucidation.

## METHODS

### Cell Culture

RAW264.7 cells (ATCC) and immortalized bone marrow-derived macrophages (iBMDM) were cultured in RPMI 1640 medium containing L-glutamine (VWR, 16750-070), 10% fetal bovine serum (FBS; Cytivia, SH30910.03), 25 mM HEPES, and 1% penicillin/streptomycin (Gibco, 15-140-122). RAW264.7 iNOS Knockout (KO) cells were generated as previously described [8,20,21]. HEK293T cells (ATCC) were cultured in DMEM medium containing high glucose, GlutaMAX™ (Thermo Scientific, 35050061) and pyruvate (Gibco, 10569044), and supplemented identically. Cell cultures were maintained at 37°C in a humidified atmosphere with 5% CO_₂_. Cells were routinely tested for mycoplasma contamination and maintained at low passage numbers. For classical activation, iBMDMs were treated with 50 ng/mL lipopolysaccharide (LPS) from *Escherichia coli* O111:B4 (Sigma-Aldrich, L3024) and 30 ng/mL recombinant mouse interferon-γ (IFNγ; R&D Systems, 485-MI-100) for the indicated durations; RAW264.7 cells were treated with 50 ng/mL LPS and 10 ng/mL IFNγ for the indicated durations.

For stable isotope labeling experiments, cells were incubated with RPMI 1640 medium containing L-glutamine media deplete of D-glucose (Gibco, 11879020) and replaced with U-^13^C-D-glucose (Cambridge Isotope Laboratories, CLM-1396-1) at formulation concentrations. 10% (v/v) Dialyzed fetal bovine serum (DFBS; Cytivia, SH30079.03) was used in place of FBS in stable isotope label containing media but otherwise supplemented with 1% (v/v) penicillin/streptomycin and 25 mM HEPES as described above.

### Primary Bone Marrow-Derived Macrophages

Primary bone marrow-derived macrophages (BMDMs) were isolated from 8–12-week-old male and female wild-type C57BL/6J mice or B6.129P2-*Nos2*^tm1Lau^/J (iNOS KO) mice (Jackson Laboratory). All animal procedures were approved by the University of Wisconsin-Madison Institutional Animal Care and Use Committee. Mice were group-housed on a 12-hour light/dark cycle at 18–23°C with 40–60% humidity and provided food and water ad libitum.

Bone marrow was flushed from femurs and tibias and pooled by genotype. Cells were differentiated into macrophages on petri dishes in RPMI 1640 medium containing L-glutamine (VWR, 16750-070), 10% FBS (Hyclone, SH30910.03), 25 mM HEPES, 1% penicillin/streptomycin (Gibco, 15-140-122), and fibroblast-conditioned medium (as a source of M-CSF). Media was replaced on days 3, 5, and 6. On day 7, differentiated BMDMs were harvested, seeded, and cultured in RPMI 1640 complete medium supplemented with 20 ng/mL recombinant mouse M-CSF (R&D Systems, 416-ML-050).

For classical activation, BMDMs were treated with 50 ng/mL LPS (E. coli O111:B4; Sigma-Aldrich, L3024) and 30 ng/mL recombinant mouse IFNγ (R&D Systems, 485-MI-100) for the indicated durations. For Nrf2 inhibition experiments, 10 µM ML385 (MedChem Express, HY-100523) was added 1 hour prior to and maintained throughout classical activation. For exogenous NO treatment, 200 µM diethylenetriamine NONOate (DETA-NONOate; Cayman Chemical, 82120) was added and maintained for the indicated durations.

### CRISPR-Cas9 based genetic knockout of *Akr1a1* in iBMDMs and RAW264.7 cells

*Akr1a1* knockout (KO) iBMDMs and *Akr1a1* KO or *Akr1a1/iNOS* double knockout (DKO) RAW264.7 cell lines were generated by CRISPR-Cas9 genome editing and method adapted from a previously described protocol [20,21]. For iBMDMs, 2 × 10□ cells were transfected via electroporation with ribonucleoprotein (RNP) complexes containing 1 µM fluorescent trans-activating CRISPR RNA (IDT, tracrRNA-ATTO550), 1 µM CRISPR RNA targeting mouse Akr1a1 (crRNA Guide #1, IDT Design ID: Mm.Cas9.Akr1a1.1.AA; or crRNA Guide #2, IDT Design ID: Mm.Cas9.Akr1a1.1.AB), and 1 µM HiFi Cas9 nuclease (IDT, 1081060) in 100 µL Nucleofector solution with supplement from Mouse Macrophage Nucleofector Kit (Lonza, VPA-1009). For RAW264.7 cells, 2 × 10□ cells were transfected via electroporation with ribonucleoprotein (RNP) complexes containing 1 µM fluorescent trans-activating CRISPR RNA (IDT, tracrRNA-ATTO550), 1 µM CRISPR RNA targeting mouse Akr1a1 (crRNA Guide, IDT Design ID: Mm.Cas9.Akr1a1.1.AA) and/or iNOS (crRNA Guide, IDT Design ID: Mm.Cas9.NOS2.1.AA), and 1 µM HiFi Cas9 nuclease (IDT, 1081060) in 100 µL Nucleofector solution V plus supplement (Lonza, VCA-1003). Electroporation was performed using program B-032 on a Nucleofector II/2b device (Lonza).

Immediately following electroporation, cells were plated in 35-mm dishes containing RPMI 1640 medium with 10% FBS without antibiotics. After 24 hours, tracrRNA-positive cells were single-cell sorted by fluorescence-activated cell sorting (FACS; BD FACSAria III) into 96-well plates containing RPMI 1640 complete medium (10% FBS, 25 mM HEPES, 1% penicillin/streptomycin). Clonal populations were expanded and screened for Akr1a1 and/or iNOS knockout by SDS-PAGE and immunoblotting with anti-Akr1a1 or anti-iNOS antibody.

### TMTpro-16plex-labeling of RAW264.7 cells

#### Cell lysis, digestion, and TMTpro labeling

RAW264.7 cell pellets (16 samples in total; ≥200 µg protein per pellet) were thawed on ice and lysed in Urea buffer (8 M urea, 100 mM Tris, pH 8.0) to an estimated concentration of 3 mg/mL. Protein concentrations were determined by A280 measurements (NanoDrop). Aliquots corresponding to 50 µg of total protein were transferred into new tubes and adjusted to a final concentration of 1 mg/mL with Urea buffer. Cysteine residues were reduced and alkylated by addition of a 10X stock solution containing TCEP (100 mM) and chloroacetamide (400 mM) in water, followed by incubation for 30 min at room temperature in the dark. The urea concentration was subsequently diluted to 1 M with 100 mM Tris, pH 8.0. Next, proteins were digested overnight at room temperature using Trypsin (sequencing-grade frozen trypsin, 0.4 µg/µL; Promega, V5113) and Lys-C (mass-spectrometry grade, 1 µg/µL in water; Wako Chemicals, 129-02541) at an enzyme-to-protein ratio of 1:50 (w/w) for each enzyme. Digestion was quenched by acidifying samples to a final concentration of 0.5% TFA. Samples were individually labeled with isobaric label following manufacturer’s protocol (Thermo Scientific, A44521). Samples were pooled by equal ratio based on labeling efficiency determined by TMTpro modified to total peptide measurements.

#### High-pH reversed-phase fractionation of TMTpro-16plex-labeled RAW264.7 peptides

After TMTpro sample pooling, 100 µg of the multiplexed peptides were subjected to a combined cleanup and high-pH reversed-phase fractionation workflow using solid-phase extraction (SPE) cartridges (1 cc, 50 mg C18 sorbent; Sep-Pak, Waters) operated on a vacuum manifold. Fractionation buffers consisted of 10 mM ammonium formate, adjusted to pH 10, and containing varying concentrations of acetonitrile (ACN). Equilibration and washing steps were performed with 1 mL of solution per step, while fractions were collected in 600 µL volumes.

#### Equilibration, sample loading, and washing

Each cartridge was first equilibrated with 100% ACN, followed by two washes with 0.2% formic acid (FA). The 100-µg peptide mixture was then loaded onto the cartridge, followed by one additional wash with 0.2% FA and one wash using a 10% ACN fractionation buffer.

#### High-pH fractionation

Peptides were eluted in a stepwise manner using 10 mM ammonium formate (pH 10) containing 18%, 20%, 22%, 24%, 26%, 28%, 30%, 32%, 34%, and 80% ACN, yielding a total of 10 fractions. The eluates were dried in a vacuum concentrator and resuspended in 0.2% FA prior to LC-MS analysis.

#### LC-MS analysis of TMTpro-16plex-peptides

For LC–MS analysis, a Vanquish Neo UHPLC system (Thermo Fisher Scientific) was coupled to an Orbitrap Ascend mass spectrometer (Thermo Fisher Scientific) equipped with a Nanospray Flex ion source. Peptide separation was performed on a 40 cm in-house–packed fused-silica column (C18, 1.7 µm, 130 Å) maintained at 50 °C using a custom-built column heater. The spray voltage was set to 2 kV, and the ion transfer tube temperature was 275 °C. TMTpro-labeled peptide fractions (500 ng total per injection) were separated at a flow rate of 300 nL/min using a 109-min active gradient from 14% to 41% solvent B (solvent A: 0.2% formic acid in water; solvent B: 80% acetonitrile with 0.2% formic acid). MS analysis was performed using an MS3–SPS–RTS (MultiNotch Synchronous Precursor Selection Real-Time Search) strategy on the Orbitrap Ascend mass spectrometer.

#### MS1 acquisition and MS2 precursor filtering

Full MS1 scans were acquired in the Orbitrap at a resolution of 60,000 over a scan range of 400–1600 m/z. The maximum injection time was 50 ms, with an automatic gain control (AGC) target of 4 × 105 (normalized AGC target = 100%). The RF lens was set to 30%. Monoisotopic peak determination (MIPS) was set to peptide, and the isolation window center was defined as the most abundant peak. Precursor charge states of 2–5 were included. Dynamic exclusion was enabled for 20 s with a ±5 ppm mass tolerance.

#### MS2 acquisition and MS3 precursor filtering

MS2 spectra were acquired in the ion trap using a 0.8 m/z isolation window and collision-induced dissociation (CID) at a normalized collision energy (NCE) of 34%, with an activation time of 10 ms and an activation Q of 0.25. Multistage activation was disabled. The ion trap scan rate was set to Turbo over a scan range of 400–1600 m/z, with a normalized AGC target of 100% and a maximum injection time of 23 ms. Real-time searching (RTS) was enabled against the UniProt Mouse Reference Proteome (UP000000589, including common contaminants; downloaded June 2024). Search parameters were as follows: enzyme = Trypsin/P; static modifications = carbamidomethyl (C, +57.0215 Da) and TMTpro16plex (Kn, +304.2071 Da); variable modifications = oxidation (M, +15.9949 Da); maximum of 1 missed cleavage and up to 2 variable modifications per peptide. Under Peak Selection and Threshold Settings, TMT SPS MS3 mode was enabled, and maximum search time was set to 80 ms. Scoring thresholds were set to: Xcorr = 1.4, dCn = 0.1, precursor mass tolerance ± 20 ppm, and charge state = 2. In the compound filter field, entries containing the keyword contam_ were rejected. Additional precursor selection filters were as follows: precursor selection range = 400–1600 m/z; precursor ion exclusion tolerance = ±10 ppm; and isobaric tag loss exclusion = TMTpro.

#### MS3 acquisition

MS3 spectra were acquired in the Orbitrap with synchronous precursor selection (SPS) enabled, selecting 10 SPS precursors using a 0.8 m/z isolation window. MS3 scans were acquired at a resolution of 45,000 (at m/z 200) over a scan range of 110–1000 m/z, with a normalized AGC target of 250% and a maximum injection time of 105 ms. High-energy collision dissociation (HCD) was applied at 55% NCE. The total cycle time (MS1 through MS3) was 3 s.

#### Proteomics Data Analysis

RAW files from LC–MS runs were processed in MaxQuant (version 2.4.2.0). Unless otherwise noted, all parameters were kept at their default settings. Group-specific parameters: In the Type field, Reporter MS3 was selected. TMTpro16plex correction factors were imported from the manufacturer’s data sheet, while all remaining settings were left at default. Global parameters: Under Sequences, the UniProt Mouse Reference Proteome (UP000000589; downloaded June 2024) was used. Under Identification, Match between runs was enabled.

Bioinformatic data analysis: Processing of MaxQuant output tables was performed in Perseus based on the proteinGroups.txt file. Entries marked as Only identified by site, Reverse, or Potential contaminant were removed. Proteins were retained only if all 16 TMTpro reporter channels contained valid (>0) values in the Reporter intensity corrected columns. These intensity values were then normalized to relative percentages, such that the total signal per channel equaled 100%. Finally, normalized intensities were log_₂_-transformed prior to downstream analysis.

### DIA analysis of Bone Marrow–Derived Macrophages

#### Cell lysis and protein digestion

Bone marrow–derived macrophage (BMDM) pellets (32 samples in total; ≥200 µg protein per pellet) were thawed on ice and lysed in Urea buffer (8 M urea, 100 mM Tris, pH 8.0) to an estimated concentration of 3 mg/mL. Lysates were briefly homogenized by probe sonication, and protein concentrations were determined by A280 measurements (NanoDrop). Aliquots corresponding to 50 µg of total protein were transferred into new tubes and adjusted to a final concentration of 1 mg/mL with Urea buffer. Cysteine residues were reduced and alkylated by addition of a 10X stock solution containing TCEP (100 mM) and chloroacetamide (400 mM) in water, followed by incubation for 30 min at room temperature in the dark. The urea concentration was subsequently diluted to 1 M with 100 mM Tris, pH 8.0. Next, proteins were digested overnight at room temperature using Trypsin (sequencing-grade frozen trypsin, 0.4 µg/µL; Promega, V5113) and Lys-C (mass-spectrometry grade, 1 µg/µL in water; Wako Chemicals, 129-02541) at an enzyme-to-protein ratio of 1:50 (w/w) for each enzyme. Digestion was quenched by acidifying samples to a final concentration of 0.5% TFA.

#### Peptide desalting

Digests were desalted using solid-phase extraction (SPE) cartridges (1 cc, 50 mg C18; Sep-Pak, Waters) operated on a vacuum manifold. Cartridges were conditioned with 100% acetonitrile (ACN), followed by two washes with 0.2% formic acid (FA). Acidified digests were then loaded onto the cartridges, washed twice with 0.2% FA, and eluted with 80% ACN containing 0.2% FA. Eluates were dried in a vacuum concentrator and resuspended in 0.2% FA for subsequent LC-MS analysis.

#### LC-MS analysis

For LC–MS analysis, a Vanquish Neo UHPLC system (Thermo Fisher Scientific) was coupled to an Orbitrap Astral mass spectrometer (Thermo Fisher Scientific) equipped with a Nanospray Flex ion source. Peptides were separated on a 40 cm in-house-packed fused-silica column (C18, 1.7 µm, 130 Å) maintained at 50 °C using a custom-built column heater. The spray voltage was set to 2 kV, and the ion-transfer-tube temperature was held at 280 °C. Peptides (250 ng total per injection) were separated at a flow rate of 300 nL/min using a 30-min active gradient from 5% to 46% solvent B (solvent A: 0.2% formic acid in water; solvent B: 80% acetonitrile with 0.2% formic acid). MS data were acquired in data-independent acquisition (DIA) mode on the Orbitrap Astral mass spectrometer (Thermo Fisher Scientific).

#### MS1 acquisition

Full MS1 scans were recorded in the Orbitrap detector at a resolution of 240,000 over a scan range of 380–980 m/z. The maximum injection time was set to 50 ms, the normalized AGC target to 250%, and the RF lens to 40%.

#### DIA acquisition

DIA scans were acquired in the Astral detector using an isolation window width of 4 m/z with no window overlap across a precursor mass range of 380–980 m/z. The DIA window type was set to Auto, and loop control was defined by time. Fragmentation was performed using higher-energy collisional dissociation (HCD) with a normalized collision energy (NCE) of 25%. The MS2 scan range was 150–2000 m/z, the injection time was 2.5 ms, and the normalized AGC target was 500%. The RF lens was maintained at 40%. The total cycle time was 0.6 s, providing rapid duty cycles for comprehensive DIA coverage.

#### Data analysis

RAW files from DIA runs were analyzed in Spectronaut (version 19.9.250512.62635; Biognosys AG) using the directDIA+ (Deep) workflow. Unless otherwise specified, all parameters were kept at their default factory settings. The UniProt Mouse Reference Proteome (UP000000589; downloaded May 2025) was used as the search database. The following non-default settings were applied: Precursor PEP cutoff = 0.01, Protein Q-value cutoff (run level) = 0.01, Protein PEP cutoff = 0.01, and Exclude single-hit proteins = True. All other parameters, including FDR control, cross-run normalization, and interference correction, followed Spectronaut’s standard BGS Factory Settings.

Protein group–level quantification data were exported from Spectronaut in pivot-transformed format. For each protein group, the number of valid PG.Quantity entries across all 32 samples was determined. Protein groups with fewer than 16 valid values were removed from further analysis. Remaining missing values were imputed in Perseus (version 2.1.3.0) using the Replace missing values from normal distribution function, with a Gaussian width of 0.3, a down shift of 1.8 standard deviations, and mode = separately for each column.

### Proteomics Data Processing and Statistical Analysis

For proteomic datasets, UniProt accession numbers were mapped to gene symbols using the UniProt.ws R package (v2.48.0) querying the mouse proteome (taxon ID: 10090). Multiple UniProt entries mapping to the same gene were collapsed by averaging quantified intensities across isoforms. BMDM protein data were back transformed from log2 (intensity_linear = 2^intensity_log2) prior to statistical analysis. This processing yielded 7,937 unique proteins in BMDM and 5,237 unique proteins in RAW264.7 cells.

Differential protein abundance was assessed using two-tailed Welch’s t-tests comparing iNOS KO + LPS/IFNγ versus WT + LPS/IFNγ groups (n=4 biological replicates per group). Statistical tests were performed on linear-scale intensity values. Log2 fold changes were calculated as log2(mean iNOS KO + LPS/IFNγ / mean WT + LPS/IFNγ). P-values were adjusted for multiple testing using the Benjamini-Hochberg false discovery rate (FDR) method, with an adjusted p-value threshold of 0.05 defining statistical significance.

### Cross-model and Cross-omic Correlation and Concordance Analysis

To assess reproducibility and identify high-confidence iNOS-regulated proteins and genes, we performed comprehensive correlation analyses across multiple datasets. All analyses compared iNOS KO + LPS/IFNγ versus WT + + LPS/IFNγ conditions unless otherwise stated. Proteins and genes were matched by gene symbol and classified based on statistical significance (adjusted p-value < 0.05) and directional agreement of log2 fold changes between datasets.

For protein-RNA correlation (RAW264.7 proteomics versus RNA-seq), proteins-genes pairs were categorized into five groups: (1) “Both sig (p-adj < 0.05) (concordant)” - significant in both datasets with log2 fold changes in the same direction; (2) “Both sig (p-adj < 0.05) (opposite)” - significant in both datasets with opposite-direction fold changes; (3 and 4) dataset-specific significance (e.g., “BMDM only” or “Protein only”); and (5) “Not significant” - not significant in either dataset. Pearson correlation coefficients were calculated between log2 fold change values for all genes with quantified data in both datasets.

A subset analysis applied stringent cutoffs to identify robust responders: genes meeting both |log2FC| > 0.5 AND adjusted p-value < 0.05 in both datasets were classified as “Both Up (>0.5)” or “Both Down (<-0.5)”. These cutoff-defined quadrants were highlighted on scatter plots with shaded regions and dashed threshold lines at ±0.5.

Triple concordant downregulated genes were identified as those showing significant downregulation (adjusted p-value < 0.05, log2FC < 0) across all three datasets (BMDM proteomics, RAW264.7 proteomics, and RAW264.7 RNA-seq), representing the highest-confidence targets of NO-dependent upregulation (i.e., genes that NO normally induces). Triple concordant genes were rank-ordered by RAW264.7 protein log2 fold change (most to least downregulated) and annotated for whether they were significantly upregulated in WT + LPS/IFNγ versus WT unstimulated (adjusted p-value < 0.05), indicating NO-inducible genes.

Results were visualized using scatter plots with Pearson correlation statistics and 95% confidence intervals. Linear regression lines with shaded confidence bands were generated using geom_smooth (method=“lm”, se=TRUE). Significance categories were color-coded by agreement type. Gene labels were added using ggrepel (v0.9.6) to prevent overlap. For model comparison with cutoffs (Figure 2C), the top 20 genes per concordant quadrant (ranked by sum of absolute log2 fold changes) were labeled, with PC2 strong contributors (defined in Principal Component Analysis section) shown in bold italic font. Significance category distributions were summarized using doughnut charts.

### Principal Component Analysis

Principal component analysis (PCA) was performed on the BMDM proteomics dataset using the prcomp function in R. All BMDM samples were included, representing combination sof genotype (WT, iNOS KO), stimulation status (unstimulated, stimulated), and DETA-NONOate treatment. Protein intensity values were log2-transformed (log2[intensity + 1]) and transposed so that samples (columns) became observations (rows). Zero-variance features were removed prior to analysis. The data matrix was centered and scaled to unit variance (scale = TRUE, center = TRUE) before computing principal components.

Variance explained by each principal component was calculated as the proportion of total variance: (PC_i standard deviation)² / Σ(all PC standard deviations)² × 100%. PC scores for each sample were extracted and annotated with experimental metadata (genotype, stimulation status, and DETA-NONOate treatment).

Genes contributing strongly to PC2 were identified using a ±2 standard deviation threshold from the mean PC2 loading value. This resulted strong positive contributors (loadings > mean + 2SD) and strong negative contributors (loadings < mean - 2SD). To examine biological relevance, mean protein expression for each condition was calculated across replicates, log2-transformed, and row-scaled (z-score normalization) for heatmap visualization. Strong PC2 contributors were clustered by correlation distance with Ward’s linkage method (ward.D2) and annotated by loading direction (positive vs. negative contributors to PC2).

### RNA-seq Data Processing and Differential Expression Analysis

Total RNA was extracted from RAW264.7 cells under matching experimental conditions (WT unstimulated, WT + LPS/IFNγ, iNOS KO unstimulated, and iNOS KO + LPS/IFNγ) with three biological replicates per group. For the Akr1a1 KO experiment, total RNA was extracted from RAW264.7 cells (WT unstimulated, WT + LPS/IFNγ, Akr1a1 KO unstimulated, and Akr1a1 KO + LPS/IFNγ) with two independent clonal replicates per group. RNA-seq was performed by Novogene using Illumina Sequencing by Synthesis technology. Quality control was performed with base recognition using CASAVA, followed by error rate assessment and GC content analysis. Data were subjected to stringent filtration: (1) removal of adaptor contamination, (2) removal of reads with >10% uncertain nucleotides (N bases), and (3) removal of reads where >50% of bases had low quality scores. After filtration, reads were aligned to the mm10 mouse reference genome using STAR, and gene-level read counts were quantified.

Gene-level read counts were imported into R (version 4.3.1) and analyzed using DESeq2 (v1.42.0). The count matrix was formatted with genes as rows and samples as columns, with sample names standardized (WT_1-3, WTstim_1-3, iNOS_1-3, iNOSstim_1-3 for the iNOS KO experiment; WT_M0a-b, WT_M1a-b, AkrKO_M0a-b, AkrKO_M1a-b for the Akr1a1 KO experiment). Genes with fewer than 10 total reads across all samples were filtered prior to analysis.

For the iNOS KO dataset, differential expression analysis was performed using the design formula ∼ condition with four condition levels (WT, WTstim, iNOS, iNOSstim). For the Akr1a1 KO dataset, an interaction design was used (∼ group, where group = genotype_treatment). The DESeq2 workflow included: (1) estimation of size factors for library depth normalization, (2) dispersion estimation using gene-wise maximum likelihood fitted to a dispersion-mean relationship, and (3) differential expression testing using the Wald test with negative binomial generalized linear models. The primary comparison for the iNOS KO experiment was iNOS KO + LPS/IFNγ versus WT + LPS/IFNγ (contrast = c(“condition”, “iNOSstim”, “WTstim”)); for the Akr1a1 KO experiment, the comparison was Akr1a1 KO + LPS/IFNγ versus WT + LPS/IFNγ (contrast = c(“group”, “AkrKO_M1”, “WT_M1”)).

P-values were adjusted for multiple testing using the Benjamini-Hochberg false discovery rate (FDR) method. For the iNOS KO comparison, genes were classified as differentially expressed if they met adjusted p-value < 0.05 AND |log_2_ fold change| > 1 (applied for volcano plot visualization and pathway enrichment). For the Akr1a1 KO comparison, only the adjusted p-value < 0.05 threshold was applied without a fold change filter, to capture the full spectrum of transcriptional changes. Gene symbols were retained from the input count matrix for downstream integration with proteomics data.

### Gene Set Definitions

ETC complex genes (Complexes I-V) were manually curated based on mitochondrial respiratory chain nomenclature. COX assembly chaperones (Cox10, Cox11, Cox15, Cox17, Cox19, Cox20) were analyzed separately from Complex IV structural subunits. Additional OXPHOS genes were obtained from KEGG (msigdbr v7.5.1, C2:CP:KEGG_LEGACY, Mus musculus). V-ATPase genes (Atp6v prefix) were distinguished from mitochondrial ATP synthase.

PC2 contributors were annotated using WikiPathways 2024 Mouse gene sets via Enrichr: (1) PC2 Negative: ETC/OXPHOS (WP295, WP1248) and Cell Cycle/IL-17A (WP413, WP5242); (2) PC2 Positive: Oxidative Stress/Glutathione (WP4466, WP412, WP164, WP730). Gene names were normalized for case-insensitive matching.

### Heatmap Generation and Normalization

Expression heatmaps were generated using pheatmap (v1.0.12). Log2-transformed intensities were row-scaled (z-score: mean=0, SD=1) to highlight relative patterns. Samples were ordered by genotype and treatment (BMDM: 32 samples including DETA conditions; RAW264.7: 16 samples). Hierarchical clustering used Pearson correlation distance with Ward.D2 linkage, except ETC heatmaps maintained sample order (cluster_cols=FALSE). Heatmaps used diverging red-blue palette (RColorBrewer RdBu) with z-scores capped at ±2.

For RAW264.7 NO-dependence annotation (Figure 2B), stimulation effects were calculated from z-scores as: WT_stim_effect = mean(WTstim) - mean(WT); iNOS_stim_effect = mean(iNOSstim) - mean(iNOS); NO_effect = WT_stim_effect - iNOS_stim_effect. Classifications: “NO-Dependent Upregulation” (WT_stim_effect > 0.4, NO_effect > 0.5), “NO-Dependent Downregulation” (WT_stim_effect <-0.4, NO_effect <-0.5), “NO-Independent” (both |stim_effects| > 0.4, |NO_effect| < 0.5), and “Under Thresholds.”

### Pathway Enrichment Analysis

Pathway enrichment was performed using Enrichr (v3.2) with hypergeometric testing against database-specific backgrounds. Analyses included: (1) PC2 contributors (WikiPathways_2024_Mouse); (2) PC2 positive TFs (ChEA_2022, ENCODE_ChEA_Consensus); (3) model comparison concordant genes (WikiPathways_2024_Mouse); (4) Akr1a1 KO DEGs (down) and combined (GO Biological Processes 2023). Results were filtered at adjusted p < 0.05 (BH-FDR). Top 15 pathways (WikiPathways) or top 5 pathways (TF analyses) were visualized with bar length =-log10 (adj-p).

### Data Visualization

All visualizations were generated in R (v4.3.1) using ggplot2 (v3.4.0) with publication-optimized themes and exported as PDF.

Volcano plots displayed log_2_FC vs.-log10(adjusted p-value) with dashed significance thresholds (p=0.05; |log2FC|=1 for iNOS KO only, no FC threshold for Akr1a1 KO). Points were colored by category (upregulated=red, downregulated=blue, not significant=gray). The top 25 genes per direction (ranked by-log10(p-adj) × |log_2_FC|) were labeled using ggrepel (v0.9.5) with italic font.

Correlation scatter plots displayed paired log_2_FC values colored by agreement category, with Pearson correlation statistics, linear regression lines (95% CI via geom_smooth), and sample sizes annotated. Model comparison plots included ±0.5 FC threshold lines and shaded quadrants, with top 20 genes per concordant quadrant labeled (PC2 contributors in bold italic).

Pathway bar plots showed top 5 (TF analyses) or top 15 (WikiPathways) terms with - log_10_(adjusted p-value) on x-axis and gene count gradient coloring. Terms were truncated to 60 characters with database prefixes removed.

PCA biplots used genotype-based colors (WT=red, iNOS KO=blue) and shapes for stimulation/DETA status, with 95% confidence ellipses (stat_ellipse, type=“norm”, level=0.95).

### Software and Statistical Computing

All proteomic and transcriptomic statistical analyses and visualizations were performed in R (version 4.3.1) using RStudio (version 2023.06.0). Core packages included: tidyverse (v2.0.0) for data manipulation, dplyr (v1.1.4) for data transformation, ggplot2 (v3.4.0) for visualization, and DESeq2 (v1.42.0) for RNA-seq differential expression analysis. Additional R packages included: pheatmap (v1.0.12) for heatmap generation, ggrepel (v0.9.5) for non-overlapping text labels, gridExtra (v2.3) for multipanel figure assembly, RColorBrewer (v1.1-3) for color palettes, msigdbr (v7.5.1) for gene set databases, enrichR (v3.2) for pathway enrichment analysis via the Enrichr web tool, UniProt.ws (v2.48.0) for protein-to-gene symbol mapping. All figures were exported as PDF files. Final figures were assembled and annotated in Adobe Illustrator 2025.

LC-MS/MS data analysis was performed using Maven (version 6.2). Immunoblots were visualized and quantified using Image Studio Lite Version 5.2 for Windows (LI-COR Biosciences). Additional experimental data not derived from proteomics or transcriptomics were graphed and analyzed in GraphPad Prism (version 10) for Windows (GraphPad Software). Final figure assembly and annotation were performed in Adobe Illustrator 2025.

Graphics were obtained from NIAID NIH BIOART: (1) Lab Mouse (bioart.niaid.nih.gov/bioart/28, 10/7/2024); (2) Petri Dish with HEK293 (bioart.niaid.nih.gov/bioart/653, 5/19/2025).

## Data Availability

The proteomics datasets have been deposited to the MassIVE repository (UC San Diego) under the accessions MSV000100329 (TMT labeling of RAW264.7 cells) and MSV000100331 (DIA analysis of bone marrow-derived macrophages). The RNA-seq dataset will be deposited to a publicly available repository. Code generated to create plots is available at: https://github.com/narp/fan_multiomic_project.git.

### Imaging of Mitochondrial Morphology

For live cell imaging, wildtype and iNOS KO RAW264.7 cells were plated at a density of 1×10^4^ cells/well onto 8-well Ibidi chambers (#1.5 glass; Ibidi, 80807) following treatment with O_2_ plasma (Diener Electronic Femto, Plasma Surface Technology) at 60 W for 3 minutes to facilitate cell adhesion. Cells were cultured in the presence and absence of LPS/IFN_γ_ for 24 hours, then incubated with MitoTracker Green (100 nM; Invitrogen, M7514) and Hoechst (1 μg/mL; Invitrogen, 33342) for one hour to label mitochondria and the cell nucleus, respectively. Chambers were then transferred to an on-stage incubator (Tokai Hit STX) set to 37°C and 5% CO_2_ for confocal imaging with a Nikon Eclipse Ti2 microscope equipped with a Crest X-light V3531 spinning disc unit and a Kinetix22 monochrome camera. Z-stacks were acquired with a Plan Apochromat 60X/1.42 NA objective with 200nm z-steps.

All image analysis was performed using FIJI/ImageJ (v2.17.0). To resolve morphology of individual mitochondria, deconvolution was performed by generating a point spread function through the ImageJ plugin DeconvolutionLab2 [59]. Mitochondrial morphological characteristics were then calculated from deconvoluted images using the FIJI/ImageJ plug-in MitoAnalyzer [60].

### Basal Oxygen Consumption Rate Measurement

To measure oxygen consumption rate (OCR) in unstimulated and LPS/IFNγ-stimulated (48 hours) wildtype and iNOS KO BMDMs with or without DETA-NONOate treatment (48 hours), 8×10^4^ BMDMs per condition were seeded in complete RPMI medium in a 96-well Agilent FluxPak tissue culture plate (Agilent, 103793-100) and treated accordingly. Media was replaced with Agilent Seahorse XF RPMI medium (Agilent, 103681-100) supplemented with 10 mM glucose, 1 mM pyruvate, and 2 mM glutamine. Cells were then incubated at 37°C for 30 minutes prior to measurement.

The Seahorse XFe96 Analyzer was calibrated with XF Calibrant solution per manufacturer’s instructions. Sensor cartridges were loaded with mitochondrial inhibitors from the Seahorse XF Cell Mito Stress Test kit (Agilent, 103015-100): 1 μM oligomycin, 0.5 μM FCCP, and 0.5 μM rotenone/antimycin A, each resuspended in assay medium. The pre-designed Mito Stress Test protocol in the Seahorse Wave Controller Software was used for OCR measurement and basal respiration calculation. OCR values were normalized to cell number using crystal violet staining as previously described [61].

### SDS-PAGE and Immunoblotting

Cells were washed with phosphate-buffered saline (Thermo Scientific, AAJ67802-K2) and lysed with RIPA Lysis Buffer (Thermo Scientific, 89901) containing Pierce™ protease inhibitors (Thermo Scientific, A32955) and phosphatase inhibitors (Thermo Scientific, A32957). Total protein content was quantified using Pierce™ BCA Protein Assay Kit (Thermo Scientific, 23227) or Pierce™ BCA Protein Assay Kit – Reducing Agent Compatible (BCA-RAC) (Thermo Scientific, 23250). Unless otherwise noted, 10 µg of total protein was loaded per well of 4-12% Bis-Tris gels (Thermo Scientific, NW04127BOX) in sample buffer containing 10% (v/v) β-mercaptoethanol (Li-COR, 928-40004). Gels were run in Bolt MES Running Buffer (Thermo Scientific, B0002) at 125 V for 65 minutes, then transferred to nitrocellulose membranes (Li-COR, 926-31092) using Bolt Transfer Buffer (Thermo Scientific, BT00061) at 12 V for 60 minutes. Membranes were blocked in 5% (w/v) non-fat dairy milk in Tris-buffered saline with 0.1% (v/v) Tween-20 (TBS-T).

Membranes were incubated with primary antibodies (1:1000 dilution in 5% [w/v] bovine serum albumin (Millipore Sigma, A7030) in TBS-T) for 4 hours at room temperature or overnight at 4°C, followed by incubation with secondary antibodies for 1 hour at room temperature or overnight at 4°C. Membranes were visualized on an Odyssey CLx imager (Li-COR) and analyzed using Image Studio Pro software (Li-COR). Primary antibodies: rabbit anti-Akr1a1 (Thermo Scientific, PA5-86685), rabbit anti-iNOS (Thermo Scientific, PIPA517106), mouse anti-α-tubulin (Cell Signaling Technologies, 3873S), rabbit anti-lipoic acid (Thermo Scientific, 43-769-5100UL), rabbit anti-DLAT (Abcam, ab66511). Secondary antibodies: goat anti-rabbit IgG 800CW (Li-COR, 926-32211), goat anti-mouse IgG 680RD (Li-COR, 926-68070). Other reagents: REVERT™ 700 Total Protein Stain (Li-COR, 926-11021).

### RT-PCR

Cells were washed with phosphate-buffered saline (Thermo Scientific, AAJ67802-K2) and lysed with RNA Stat60 (Fisher Scientific, NC9489785) with subsequent RNA isolation performed per manufacture’s protocol. Chloroform (Sigma Aldrich, 650498) was used for phase separation. RNA was precipitated with isopropanol (Thermo Scientific, AC327272500) and washed with 75% [v/v] molecular-grade ethanol (VWR, 71006-012). RNA pellets were resuspended in molecular-grade water and quantified with a NanoDrop 2000c Spectrophotomer (Thermo Scientific). cDNA was synthesized using the High-Capacity cDNA Reverse Transcription Kit (Thermo Scientific, 4368813). Quantitative PCR reactions were performed with 10 ng of cDNA per reaction using PowerUp™ SYBR™ Green Master Mix (Thermo Scientific, A25777) on an Applied Biosystems QuantStudio 7 Pro Real-Time PCR System (Thermo Scientific). Gene expression was analyzed using the ΔΔCt method with *Hnrpab* as the reference gene. The following target primers were used: *Akr1a1*, <u>Forward</u>: 5’-CTGATGCAGTCCTGCTTGA-3’, <u>Reverse</u>: 5’-GGATGCAGATCACTTTCCGCT-3’; *Hmox1*, <u>Forward</u>: 5’- AAGCCGAGAATGCTGAGTTCA-3’, <u>Reverse</u>: 5’-GCCGTGTAGATATGGTACAAGGA-3’; *Hnrpab*, <u>Forward</u>: 5’-AGGACGCGGGAAAAATGTTC-3’, <u>Reverse</u>: 5’- CAGTCAACAACCTCTCCAAACT-3’; *IL6*, Forward: 5’-CCAGAAACCGCTATGAAGTTCCT-3’, Reverse: 5’-CACCAGCATCAGTCCCAAGA-3’.

### Nitrite Concentrations

Nitrite concentrations in conditioned media were measured using the Griess Reagent System (Promega, G2930) according to the manufacturer’s instructions. Briefly, 50 µL of conditioned media was combined with sulfanilamide and N-1-napthylethylenediamine dihydrochloride solutions in a 96-well plate. Absorbance was measured at 535 nm within 30 minutes using an Epoch 2 microplate reader. Nitrite concentrations were calculated from a standard curve generated using sodium nitrite standards ranging from 0 to 1000 µM.

### PDHC Activity Assay from Cell Lysate

PDH complex activity was measured in lysates from unstimulated or LPS/IFNγ-stimulated wildtype and *Akr1a1* knockout iBMDM clones using the Pyruvate Dehydrogenase Enzyme Activity Microplate Assay Kit (Abcam, ab109902) according to the manufacturer’s instructions. Briefly, cells were lysed and protein concentrations were determined by Pierce™ BCA Protein Assay Kit (Thermo Scientific, 23227). Equal amounts of protein were loaded into the immunocapture plate and incubated for 3 hours at room temperature. Following washing, PDH activity was measured by monitoring NADH production at 450 nm using BioTek Epoch2 microplate reader. Mean velocity within the linear range was normalized to blank and expressed as relative activity to unstimulated wildtype lysate. Data were analyzed using Gen5 TS v.209 software (BioTek Instruments, Inc.).

### FLAG-tagged AKR1A1 protein expression and purification

The lentiviral-based protein expression and FLAG-tagged protein purification protocol was adapted from a previously described method [8].To generate HEK293T cells (ATCC, CRL-3216) stably expressing FLAG-tagged Akr1a1 or FLAG-tagged Akr1a1(K127A) mutant, lentiviral particles were produced by transfecting cells with FuGENE 6 Transfection Reagent (Promega, E2691) and Opti-MEM (Thermo Scientific, 31985070) containing psPAX2 packaging plasmid (Addgene, plasmid no. 12260), pMD2.G envelope plasmid (Addgene, plasmid no. 12259), and pLV[Exp]-Puro-CMV-Akr1a1[NM_021473.3]/3xFLAG (VectorBuilder, VectorID: VB250429-1427ssh) or pLV[Exp]-Puro-CMV-Akr1a1_K127A[NM_021473.3]/3xFLAG (VectorBuilder, VectorID: VB250429-1440bbq). Viral particles were added to cells in DMEM containing 8 μg/mL polybrene (Sigma-Aldrich, TR-1003) for 24 hours. Transduced cells were selected with 5 μg/mL puromycin (Sigma-Aldrich, P4512).

For protein purification, cells were washed and lysed in lysis buffer (20 mM Tris-HCl pH 7.4, 1% Triton X-100, 100 mM NaCl, 5 mM MgCl₂, 1 mM dithiothreitol (DTT) (Thermo Scientific, AAJ1539706)) containing Pierce™ protease inhibitors (Thermo Scientific, A32955) and phosphatase inhibitors (Thermo Scientific, A32957). Total protein was quantified using Pierce BCA Protein Assay Kit – Reducing Agent Compatible (BCA-RAC) (Thermo Scientific, 23250). Then, 2 mg of total protein was added to pre-washed anti-FLAG M2 resin (Sigma-Aldrich, A2220) and rotated end-over-end for 3 hours at 4°C. The resin was washed four times with lysis buffer and centrifuged at 2,000 × g for 1 minute at 4°C between washes. Following the final wash, the resin was washed once with 1 mL final wash buffer (20 mM Tris-HCl pH 7.4, 100 mM NaCl, 5 mM MgCl₂) and centrifuged at 2,000 × g for 1 minute at 4°C. Bound protein was eluted by adding 7 µg FLAG peptide in final wash buffer and rotating at 1,200 rpm for 30 minutes at 25°C. The eluate was collected by centrifugation at 1,000 × g for 4 minutes at 4°C through a chromatography column (Bio-Rad Laboratories, 7326204).

The eluted protein was transferred to a 30-kDa MWCO filter (Sigma-Aldrich, UFC503024) and buffer-exchanged into storage buffer (40 mM Tris-HCl pH 7.5, 100 mM NaCl, 2 mM DTT) by repeated concentration (10 cycles of adding 300 µL storage buffer and centrifuging at 10,000 × g for 5 minutes at 4°C). Glycerol was added to a final concentration of 15% (v/v). Protein concentration was determined by NanoDrop 2000c Spectrophotometer using an extinction coefficient and molecular weight for WT and mutant, respectively: 60,850 M⁻¹cm⁻¹ and 39,299.28 Da, 60.850 M⁻¹cm⁻¹ and 392423.18 Da. Protein purity was confirmed by SDS-PAGE with Coomassie staining (Figure 4E).

### In vitro purified AKR1A1 co-incubation PDHC activity assays

PDHC activity assay was adapted to include Akr1a1 co-incubation from a previously described protocols [20,21]. Purified porcine PDHC (Sigma Alrich, P7032-10UN) (0.2 units/mg) was incubated at room temperature for 3 hours with indicated combinations of PAPA-NONOate (600 µM), CoA (200 µM), NADH (200 µM), and NADPH (200 µM) with or without 250 nmol of purified AKR1A1-Flag or AKR1A1[K127A]-Flag in 20 mM sodium phosphate buffer (pH 7.2). After incubation, the protein mixture was diluted 1:20 in assay solution containing thiamine pyrophosphate (100 µM), CoA (2 mM), pyruvate (2 mM), and NAD+ (10 mM) in 20 mM sodium phosphate buffer (pH 7.2). PDHC activity was quantified by the rate of NADH production as measured by the increasing NADH absorbance at 340 nm over time using a BioTek Epoch2 microplate reader. Absorbance was measured continuously, and the mean velocity was determined from the linear portion of the curve. Data were analyzed using Gen5 TS v.209 software (BioTek Instruments, Inc.).

### Metabolite extraction and LC-MS analysis

To measure intracellular metabolites, cells were washed twice with D-PBS and metabolites were extracted with cold LC-MS grade 80:20 (v/v) methanol:water (Thermo Scientific, A4564, W64). The resulting pellet was extracted twice then pooled in a tube and dried under a nitrogen stream. Dried samples were resuspended in LC-MS grade H_2_O. Samples were analyzed using a Q Exactive Orbitrap Mass Spectrometer (Thermo Fisher Scientific) coupled to a Vanquish Horizon UHPLC System. Xcalibur 4.0 was used for data acquisition. Samples were separated on a 100L×L2.1Lmm, 1.7LμM ACQUITY UPLC BEH C18 Column (Waters) with a gradient of solvent A (97:3 (v/v) water:methanol, 10 mM TBA (cat. no. 90781, Sigma-Aldrich)), 9 mM acetic acid (pH 8.2, cat. no. A11350, Thermo Fisher Scientific) and solvent B (100% methanol). The gradient was: 0 min, 5% B; 2.5 min, 5% B; 17 min, 95% B; 21 min, 95% B; 21.5 min, 5% B. The flow rate was 0.2LmlLmin^−1^ with a 30 °C column temperature. Data were collected in full scan negative mode at a resolution of 70 K. Settings for the ion source were: 10 auxiliary gas flow rate, 35 sheath gas flow rate, 2 sweep gas flow rate, 3.2LkV spray voltage, 320 °C capillary temperature and 300 °C heater temperature. Reported metabolites were identified based on exact *m/z* and retention times determined using standards. Data were analyzed with MAVEN (v.2011.6.17) [62,63]. Metabolite levels were normalized to total protein content quantified using Pierce™ BCA Protein Assay Kit (Thermo Scientific, 23227).

## ELISA

To measure cytokine release, 0- or 48-hour spent medium from RAW264.7 cells was collected and spun at 500 x *g* for 5Lminutes. The IL-6 concentration in the supernatant was measured using DuoSet ELISA Kits (R&D Systems, DY406) according to the manufacturer’s instructions. The levels of released IL-6 were normalized to the total cellular protein content in the culture plate.

## ACKNOWLEDGEMENTS

The authors thank J.D. Sauer at the University of Wisconsin School of Medicine and Public Health for the immortalized bone marrow derived macrophages. The authors thank the University of Wisconsin Carbone Cancer Center Flow Cytometry Laboratory, supported by P30 CA014520, for use of its facilities and services.

---

**Supplemental Figure 1.**
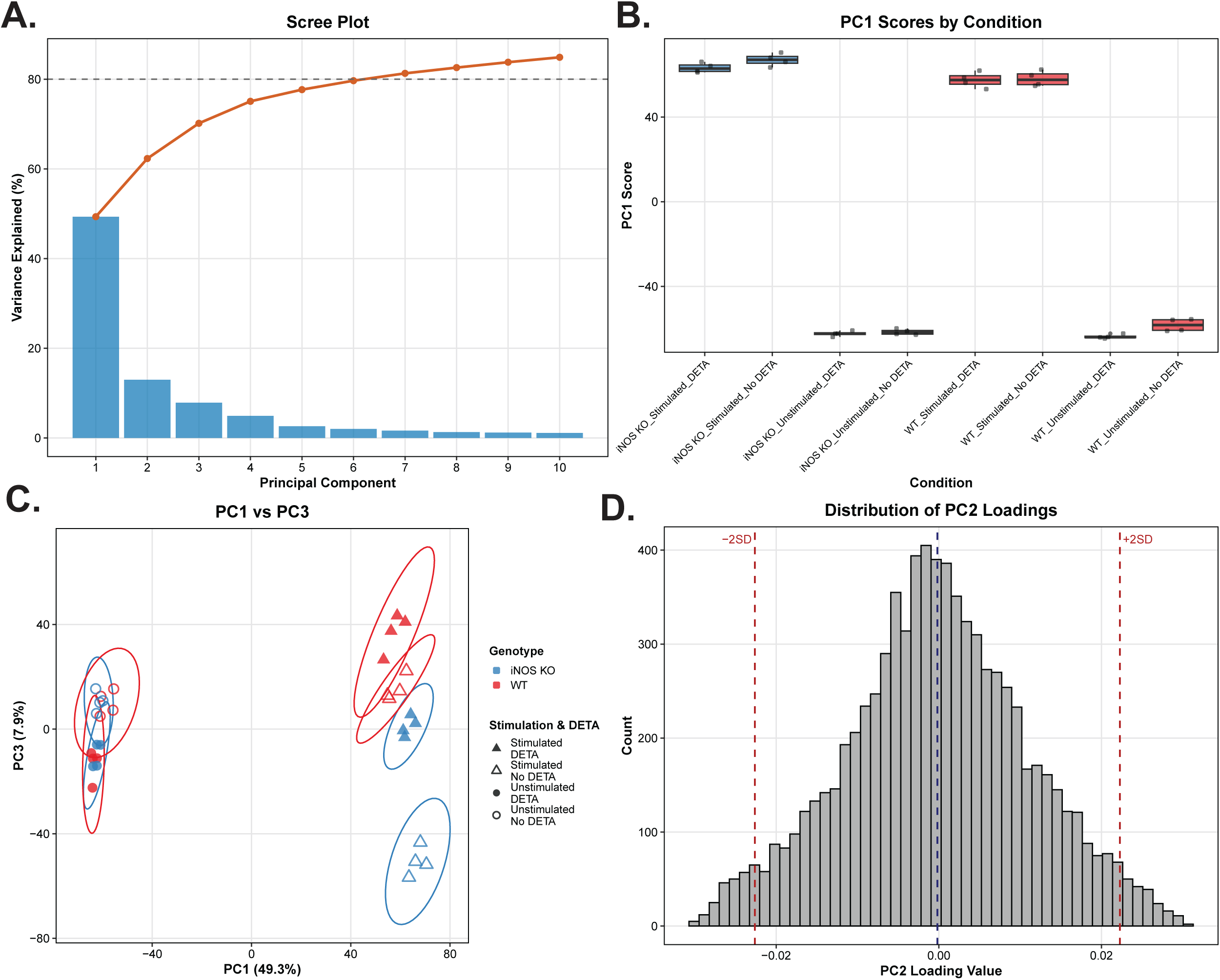
Principal component analysis (PCA) of BMDM proteomic data reveals NO•-dependent protein signatures. Principal component analysis (PCA) was performed on log_₂_-transformed protein intensities from BMDM cultures across all experimental conditions (WT and iNOS KO, unstimulated and stimulated with LPS/IFNγ for 48 hours, +/- DETA-NONOate; n=4 biological replicates per condition, 8 conditions total). **(A)** Scree plot showing variance explained by the first 10 principal components. **(B)** Distribution of PC1 scores across experimental conditions. Box plots show median, interquartile range (box), 1.5x interquartile range (whiskers), and individual replicates (points). **(C)** PCA map along PC1 and PC3. Ellipses represent 95% confidence intervals for each of the 8 unique condition groups. **(D)** Distribution of PC2 loadings across all proteins (n=7,937). Proteins with loadings >2 standard deviations from the mean were classified as strong PC2 contributors (n=209 positive contributors; n=223 negative contributors). Dashed vertical lines indicate ±2SD thresholds.

**Supplemental Figure 2.**
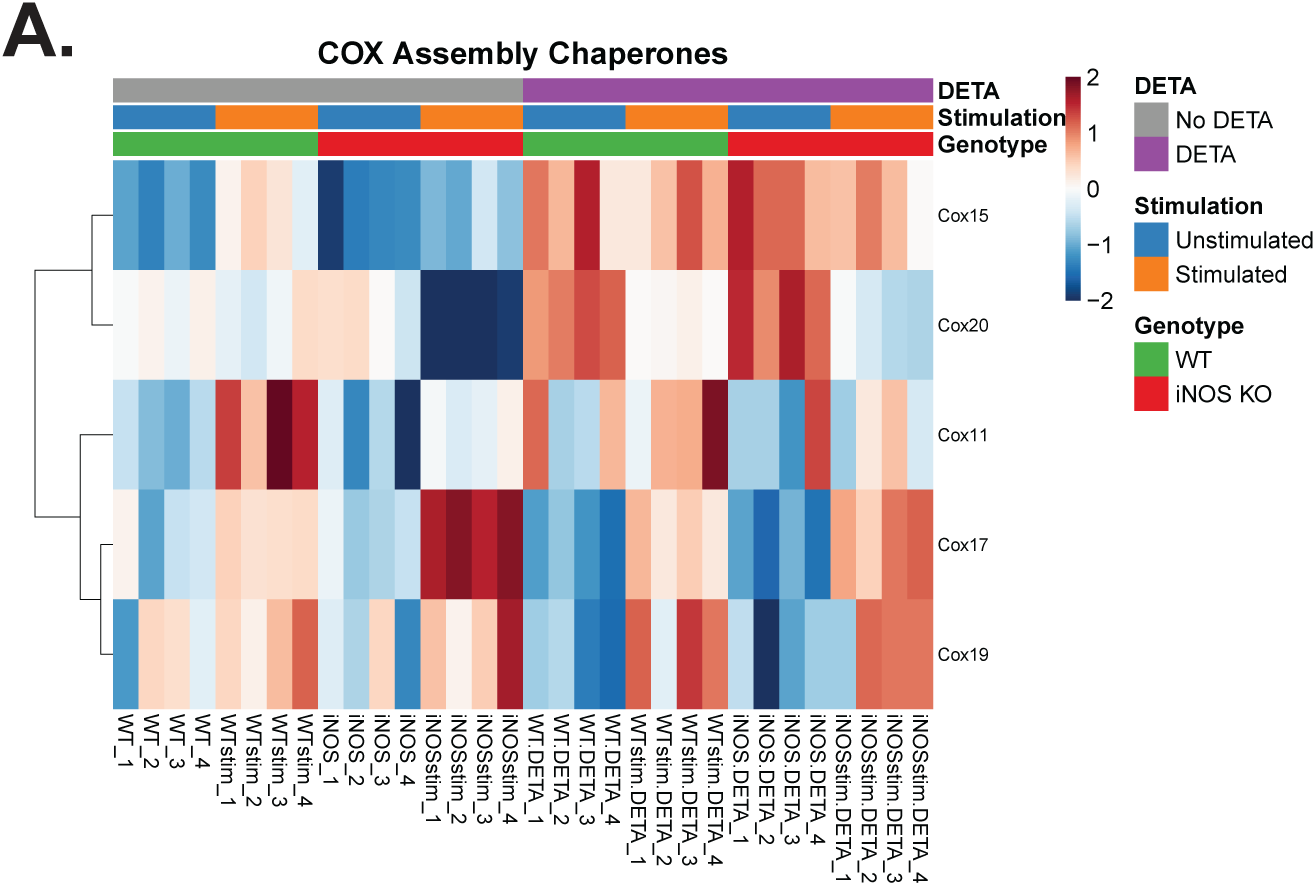
NO• does not drive COX Assembly Chaperone remodeling. **(A)** Heatmap showing changes in COX Assembly Chaperone proteins (Cox10, Cox11, Cox15, Cox17, Cox19, Cox20) in BMDM across eight experimental conditions: unstimulated (unstim) or stimulated (stim) with LPS/IFNγ for 48 hours in WT or iNOS KO genotypes; +/- DETA-NONOate (DETA). Colors represent row-wise Z-score normalized protein abundance (blue = decreased, red = increased relative to row mean). Rows (proteins) clustered by Pearson correlation; columns ordered by experimental condition. n = 4 biological replicates per condition (individual replicates designated by “_1” through “_4” suffix).

**Supplemental Figure 3.**
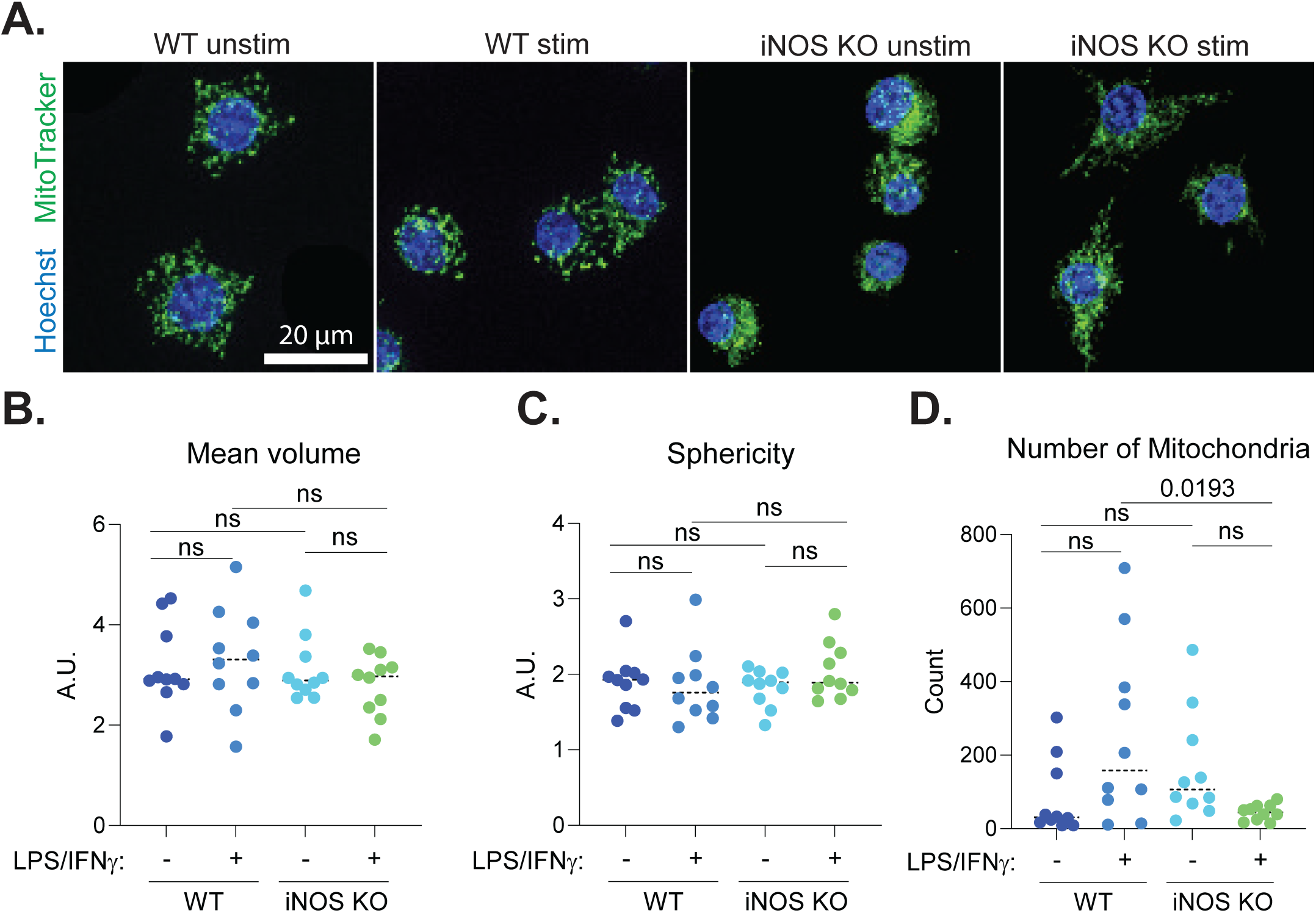
Mitochondrial morphology quantified in stimulated or unstimulated WT and iNOS KO RAW264.7 cells. **(A)** Representative live cell confocal images of wildtype (WT) and iNOS knockout (KO) RAW264.7 cells with or without LPS/IFNγ stimulation for 24 hours. Nucleus (blue, Hoechst), mitochondria (green, MitoTracker Green). Scale bar = 20 μm. **(B-D)** Quantification of mean mitochondrial volume (B), sphericity (C), and number per cell (D) from MitoTracker-labeled cells analyzed with MitoAnalyzer. Data represent mean +/- standard deviation; n = 10 cells from 2 independent experiments. Statistical comparisons by one-way ANOVA with Tukey’s post hoc test for multiple comparisons; exact p-values shown; ns = not significant.

**Supplemental Figure 4.**
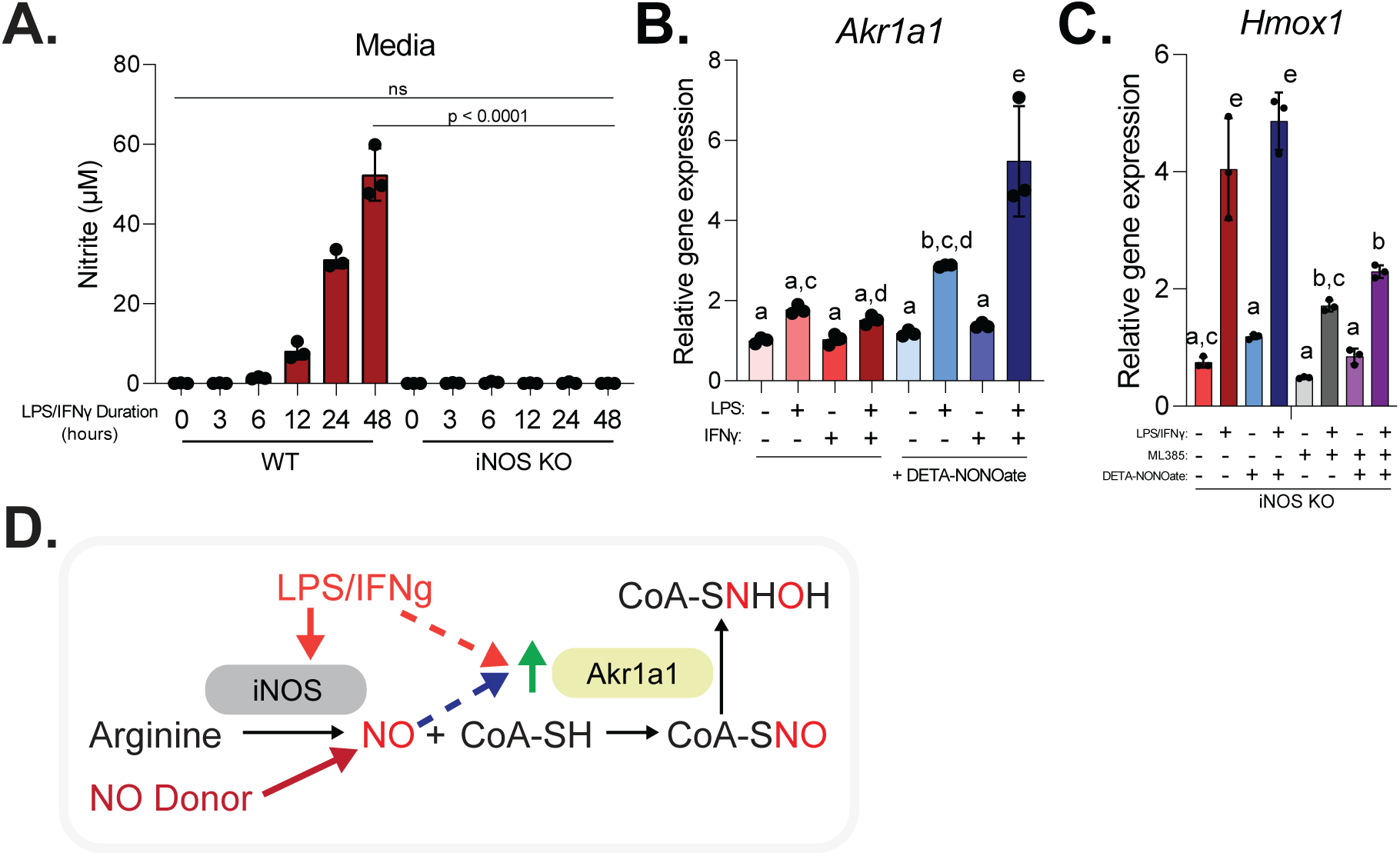
The upregulation of AKR1A1 requires both activation signal and NO. **(A)** Nitrite accumulation in culture media from cells in (Figure 5B) measured by Griess assay. Data represents mean +/- standard deviation (SD), n = 3 biological replicates. **(B)** Relative mRNA expression of *Akr1a1* in iNOS KO BMDMs treated with all combinations of LPS, IFNγ, and DETA-NONOate (200 μM) for 48 hours. Expression normalized to *Hnrpab* reference gene using ΔΔCt method. Data represents mean +/- SD, n = 3 biological replicates. **(C)** Relative mRNA expression of *Hmox1* (J), a canonical Nrf2 target, in iNOS KO BMDMs stimulated with LPS/IFNγ +/- DETA-NONOate +/- ML385 (Nrf2 inhibitor, 10 μM) for 48 hours. Expression normalized to *Hnrpab*. Data represents mean +/- SD, n = 3 biological replicates. **(D)** Schematic model depicting co-regulation of Akr1a1 by classical activation and NO signals. <u>Statistics</u>: For A-C, statistical comparisons were performed using one-way ANOVA with Tukey’s post hoc test for multiple comparisons. Bars with different letters indicate statistically significant differences (p < 0.05). ns = not significant.

**Supplemental Figure 5.**
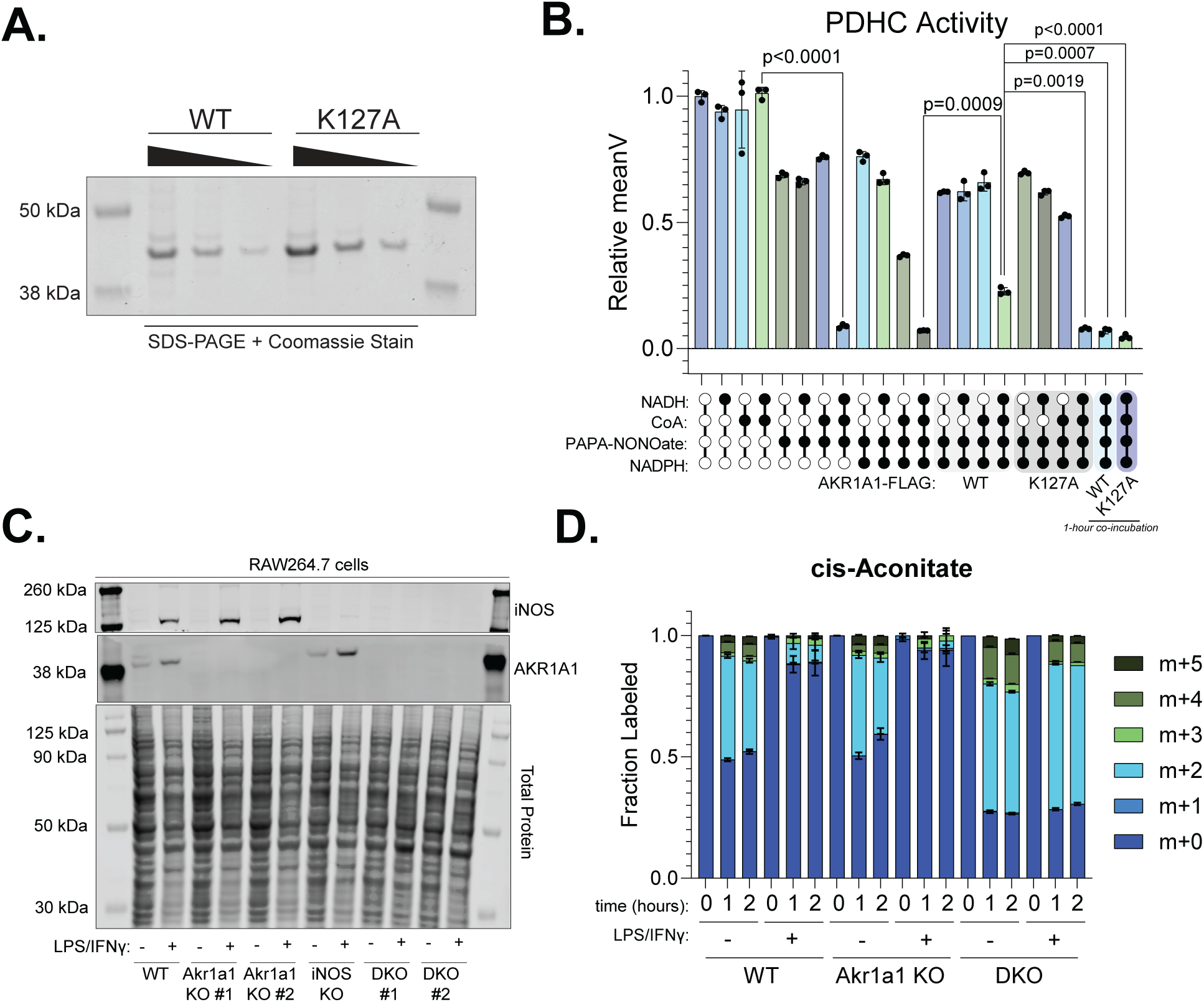
AKR1A1 regulates TCA cycle by tempering the effect of NO•. **(A**) Coomassie-stained SDS-PAGE gel showing purified recombinant AKR1A1-FLAG proteins. WT and catalytically inactive K127A mutant AKR1A1-FLAG were expressed in HEK293T cells and purified via anti-FLAG affinity chromatography. Serial dilutions (left to right) demonstrate purity. Expected molecular weight: ∼40 kDa. **(B)** Temporal requirement for Akr1a1 protection. Same conditions as (Figure 6E) but with all conditions represented and two additional: Akr1a1-FLAG (WT or K127A) added after 2-hour pre-incubation with NADH, NADPH, CoA, and PAPA-NONOate versus Akr1a1 added at time zero. Activity normalized to protein only control. Data represents mean +/- SD, n = 3 independent reactions. **(C)** Immunoblot showing iNOS and AKR1A1 protein abundance for WT, *Akr1a1* KO, iNOS KO, and *Akr1a1/iNOS* DKO clones (#1 and #2 indicate independent single clones) in RAW264.7 cells. All unstimulated or stimulated with LPS/IFNγ for 48-hours. Total protein stain shown. **(D)** Isotopologue distribution of cis-Aconitate from conditions described in Figure 6F. <u>Statistics</u>: For panels B, statistical comparisons were performed using one-way ANOVA with Tukey’s post hoc test for multiple comparisons. Exact p-values reported.

**Supplemental Figure 6.**
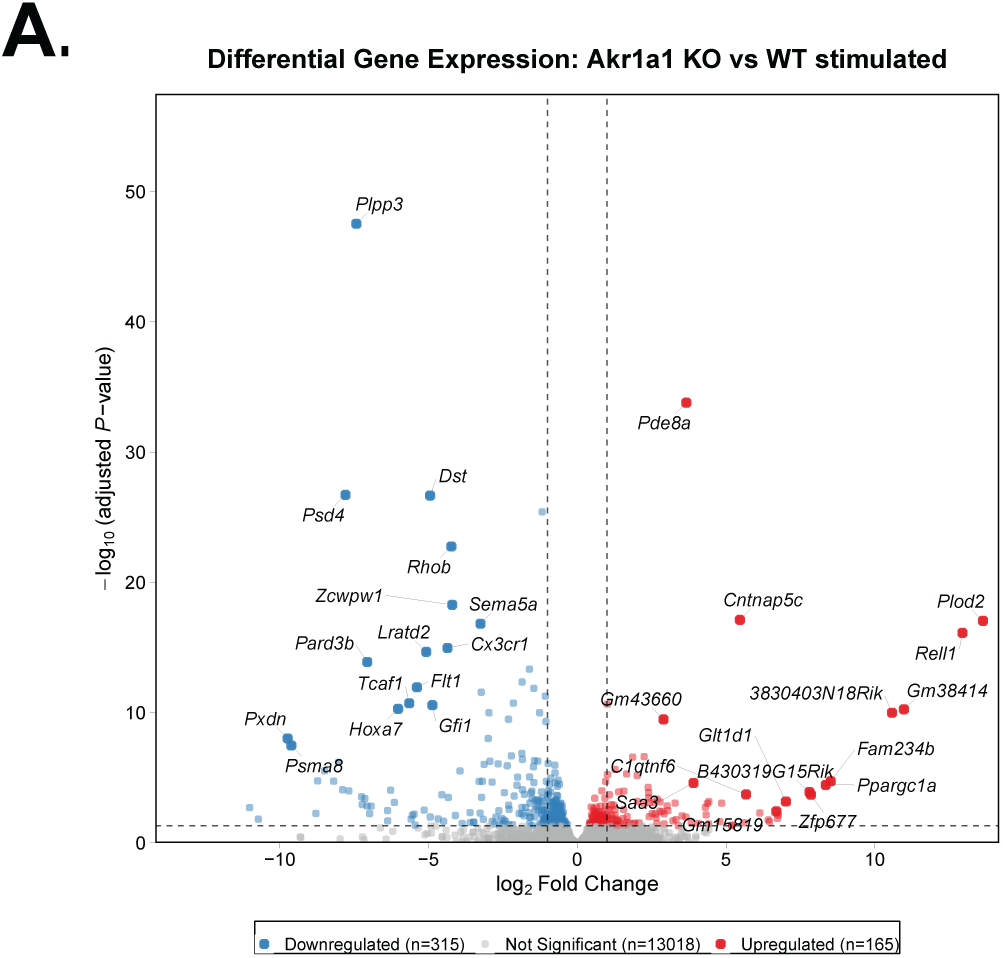
A*k*r1a1-dependent transcriptome changes observed in stimulated RAW264.7 cells. **(A)** Volcano plot showing differentially expressed genes in RAW264.7 cells comparing stimulated Akr1a1 KO versus WT (LPS/IFNγ, 48 hours; n = 2 independent clonal replicates per genotype). Genes meeting significance threshold (p-adj < 0.05) are colored by direction of change: upregulated in Akr1a1 KO (red), downregulated in Akr1a1 KO (blue), not significant (grey). Dashed lines indicate thresholds at p-adj = 0.05 (horizontal) and |log2FC| = 1 (vertical) for reference. Top 25 genes per direction (ranked by-log10(p-adj) × |log2FC|) are labeled.

